# An organoid model of the menstrual cycle reveals the role of the luminal epithelium in regeneration of the human endometrium

**DOI:** 10.1101/2025.07.03.663000

**Authors:** Konstantina Nikolakopoulou, Weand Ybañez, Lhéanna Klaeylé, Lisa Frugoli, Hans-Rudolf Hotz, Charlotte Soneson, Margherita Yayoi Turco

**Affiliations:** Friedrich Miescher Institute for Biomedical Research, Fabrikstrasse 24, 4056 Basel, Switzerland; SIB Swiss Institute of Bioinformatics, Basel, Switzerland

## Abstract

Menstruation is an unusual physiological process whereby the human endometrium undergoes cyclical shedding yet scarless regeneration. Despite its pivotal role in reproductive health, the cellular states and interactions that coordinate this process are incompletely defined. Here, we establish an *in vitro* menstrual cycle (IVMC) protocol using human endometrial organoids that faithfully recapitulates the epithelium across the key phases of the menstrual cycle including differentiation, hormonal withdrawal, breakdown and regeneration. This IVMC protocol enables intricate study of the early phases of the cycle, which remain difficult to access *in vivo.* Using this system, we define transcriptional transitions in ciliated and non-ciliated epithelial cells during regeneration, revealing immediate stress-responsive states followed by wound-responsive ones that precede proliferation. We demonstrate that, in response to breakdown, organoids acquire a transcriptomic signature resembling the *in vivo* luminal epithelium in the menstrual and proliferative phases of the cycle. Luminal epithelial cells are crucial for this re-epithelialization and express factors such as *WNT7A* which we show to be required for long-term epithelial maintenance. Moreover, cell–cell communication analyses identify luminal epithelium as a signaling hub, interacting with endothelial and immune cells via pathways including CXCL8, consistent with promotion of angiogenesis and immune recruitment during the regenerative window. Taken together, our study establishes a physiologically relevant paradigm to dissect epithelial renewal and cell-cell interactions during menstruation with potential to extend towards modelling common and distressing conditions such as menstrual disorders and endometriosis.

## Introduction

Menstruation is a remarkable physiological process where the uterine mucosa, the endometrium, sheds and regenerates in a scarless manner. The endometrium is the site of implantation and growth of the conceptus and undergoes cyclical functional and morphological remodelling during the menstrual cycle in a process that is tightly regulated by the ovarian hormones estrogen and progesterone^1^ (Fig.1a). During each cycle, the endometrium faces a binary fate. When implantation occurs, the upper functional layer is maintained and differentiates into the decidual lining required to support pregnancy. In the absence of implantation, with the involution of the corpus luteum, there is a sharp decline in progesterone that initiates menstruation. This is tightly coordinated and involves local hypoxia, infiltration of leukocytes and activation of matrix metalloproteases by stromal cells, which promote degradation of extracellular matrix and tissue sloughing^2^. As breakdown proceeds, rapid re-epithelialization of the denuded surface is initiated followed by restoration of the underlying stroma and vascular compartments to fully regenerate the functional layer^3^. This cyclical breakdown and scarless regeneration of the endometrium, is limited to few species and correlates with the extent of intrusion of the blastocyst into the uterine mucosa which is most extreme in humans^4^.

**Figure 1:**
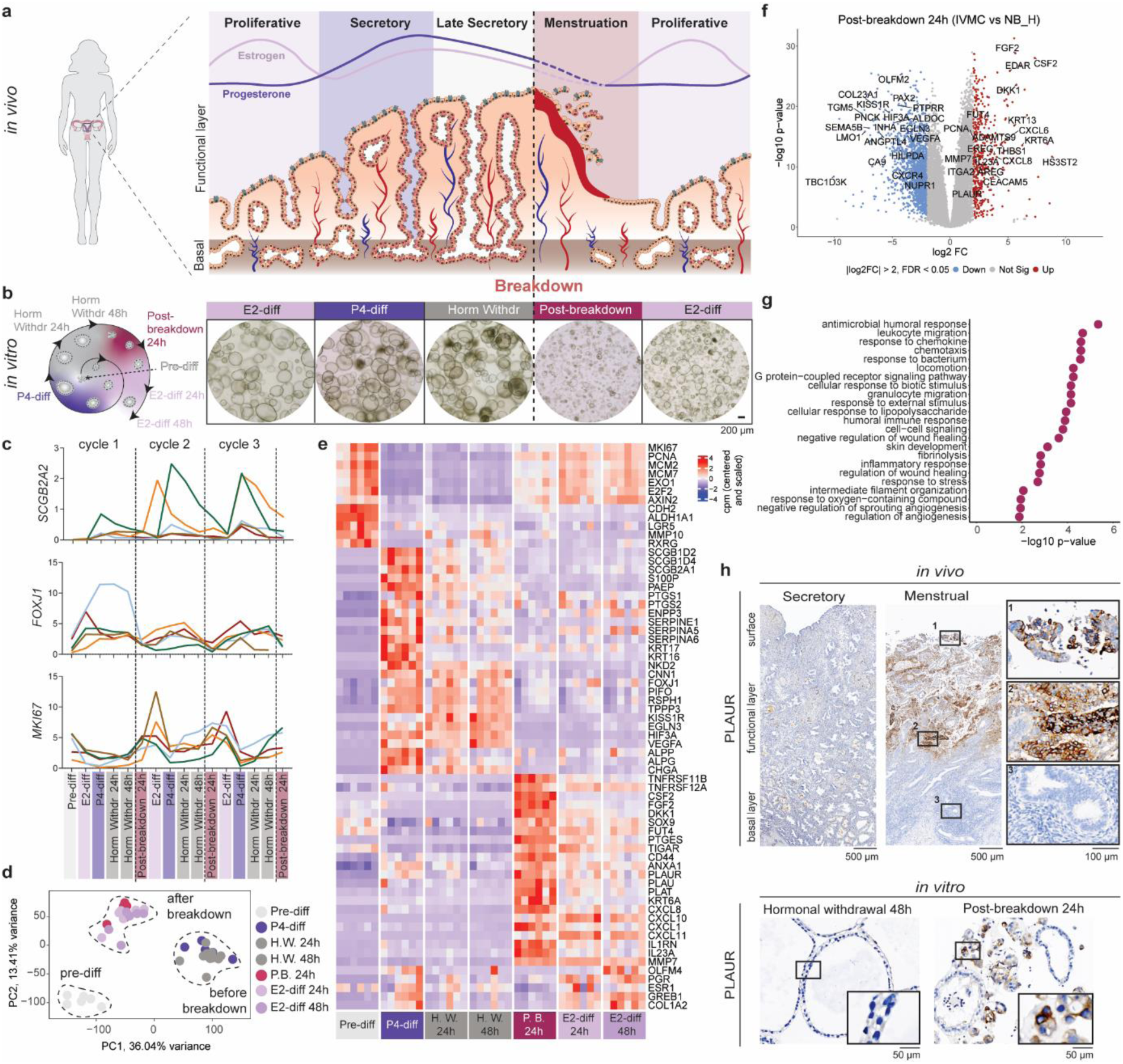
An *in vitro* menstrual cycle protocol (IVMC) recapitulates the dynamics of the endometrial epithelium. (**a**) Schematic of the morphological changes of the human endometrium and fluctuation of ovarian hormones, estrogen and progesterone, during different phases of the menstrual cycle. (**b**) Schematic of the IVMC protocol starting from EOs growing from single cells (left) and representative brightfield images (right). (**c**) qRT-PCR analysis of *SCGB2A2* (secretory marker), *FOXJ1* (ciliated marker), and *MKI67* (proliferative marker) expression from EOs undergoing continuous cycles of the IVMC protocol, normalised to housekeeping genes (n=5 independent EO lines). (**d**) PCA of batch corrected samples in the IVMC protocol analysed by bulk RNA sequencing, coloured by timepoint (n=6 independent EO lines). (**e**) Heatmap depicting centered and scaled counts per million (cpm) of selected genes across batch corrected samples in the IVMC protocol. (**f**) Volcano plot highlighting selected DEGs of EOs in Post-breakdown 24h phase of the IVMC protocol compared to control EOs (NB_H). (**g**) Biological processes enriched in Post-breakdown 24h phase of the IVMC protocol using genes upregulated in (f). (**h**) Representative IHC image for PLAUR in sections from endometrial tissue of the secretory and menstrual phases (n=3 donors for each phase) and EOs at Horm Withdr 48h and Post-breakdown 24h (n=4 independent EO lines). Boxed areas of EO sections are three times magnified. Abbreviations: IVMC; *in vitro* menstrual cycle, Pre-diff; Pre-differentiation, P4-diff; Progesterone differentiation, Horm Withdr; Hormonal Withdrawal, E2-diff; Estrogen differentiation, NB_H; No Breakdown_Hormones.

Despite the central importance of menstruation throughout reproductive life, the cellular states involved, and their coordination remain poorly understood. The epithelial compartment is a key component of the endometrium and is thought to play a central role in its cyclical remodelling. The basal portions of the glands are not shed during menstruation and have been considered the main source of regenerative cells for restoring the epithelium^3,5^. However, the upper portions of the endometrial glands could also mediate both tissue breakdown and regeneration with the luminal cells contributing to re-epithelialization as well as potentially serving as another reservoir of stem/progenitor cells^6–9^. How these different epithelial cell states emerge and transition during the regenerative phase of the menstrual cycle remains to be defined.

Studying this complex cyclical process has been challenging due to the lack of physiologically relevant model systems^10,11^. Here we use human endometrial organoids (EOs), that can be maintained long-term in culture and which recapitulate the epithelial compartment and its responsiveness to hormones^12–14^. Using EOs, we established an *in vitro* menstrual cycle (IVMC) protocol that recapitulates key features of the cycle, including epithelial differentiation, hormonal withdrawal, mechanical breakdown and epithelial renewal. The IVMC protocol enables detailed investigation of epithelial dynamics over multiple cycles and reveals a critical role of luminal epithelial cells in endometrial regeneration. Overall, this tractable platform allows further investigation of the molecular mechanisms and cell-cell interactions governing epithelial regeneration in physiological conditions and in disease states such as abnormal menstrual bleeding and endometriosis.

## Results

### An *in vitro* menstrual cycle protocol (IVMC) recapitulates the dynamics of the endometrial epithelium

To study the cyclical changes at endometrial breakdown and regeneration, the hallmarks of the menstrual window, we developed an *in vitro* menstrual cycle (IVMC) protocol that simulates these critical processes using EOs (Fig.1b). EOs respond to ovarian hormones and differentiate into the two main epithelial cell types: ciliated and secretory cells^12–14^. Established EO cultures were dissociated into single cells and cultured until organoid formation occurred (Pre-differentiation; Pre-diff). This timepoint marks the beginning of the first IVMC cycle representing the state of organoids in standard culture conditions under expansion. EOs were then cultured with estrogen (E2) to mimic the proliferative phase, followed by the addition of progesterone (P4), prolactin (PRL), and cyclic adenosine monophosphate (cAMP) to simulate the secretory phase (P4-diff), as previously described^12^. To mimic the rapid decline in ovarian hormones occurring with the demise of the corpus luteum in the absence of embryonic implantation, all hormonal supplements and cAMP were withdrawn. EOs were collected 24 and 48 hours after hormonal withdrawal (Horm Withdr 24h; Horm Withdr 48h). The key mediators of menstrual breakdown *in vivo* are matrix metalloproteinases produced by stromal and immune cells^15,16^. In their absence, EOs were mechanically disrupted to mimic epithelial breakdown. To simulate the regenerative phase, the resulting fragmented EOs were replated and first maintained without hormones for 24 hours (Post-breakdown 24h) before they get exposed to E2 for 24 and 48 hours (E2-diff 24h; E2-diff 48h) (Fig.1b). This cyclical process can be repeated multiple times. Quantitative real-time PCR (qRT-PCR) analysis of secretory (*SCGB2A2*) and ciliated (*FOXJ1*) markers across three consecutive cycles confirms cyclical expression patterns corresponding to hormonal exposure, whereas the proliferative marker (*MKI67*) demonstrates an inverse pattern (Fig.1c).

To validate that our IVMC protocol recapitulates the epithelial changes throughout the menstrual cycle *in vivo*, bulk RNA sequencing of six independent EO lines was performed at defined time points. Control conditions included (i) EOs not broken down but treated with hormones (No Breakdown Hormones; NB_H), (ii) EOs broken down but not treated with hormones (Breakdown No Hormones; B_NH), and (iii) EOs neither broken down nor treated with hormones (No Breakdown No Hormones; NB_NH) (Extended Data Fig.1a). Mechanical disruption is the major source of transcriptomic variance within all conditions and timepoints, with hormonal treatment contributing to secondary segregation, as shown by principal component analysis (PCA) (Extended Data Fig.1b, c). Organoids in the IVMC protocol show distinct clustering corresponding to the differentiation status and stages of breakdown (Fig.1d). Comparison of each time point of the IVMC protocol to all other timepoints reveals distinct gene expression patterns associated with each phase (Extended Data Fig.1d, Supplementary Tables).

We further validated our protocol by interrogating differentially expressed genes (DEGs) across different timepoints of the IVMC, in combination with known markers that were confirmed by immunohistochemistry (IHC) (Fig.1e). EOs at the Pre-diff timepoint have a proliferative signature *(MKI67*, *MCM7*, *PCNA*), representing their state in standard culture conditions for expansion (Fig.1e, Extended Data Fig.2a, b). Putative markers of endometrial stem/progenitor cells, including *ALDH1A1, CDH2, AXIN2*, are also expressed in Pre-diff EOs^17–19^ (Fig.1e, Extended Data Fig.2a, c). Differential expression analysis (DEA) comparing hormonally-treated and control EOs (B_NH) at the P4-diff timepoint shows upregulation of progesterone-regulated genes (*PTGS2, ENPP3, SCGB1D4*)^20^ (Fig.1e, Extended Data Fig.3a, Supplementary Tables). Genes associated with the secretory lineage (*PAEP*, *S100P, SCGB2A1, SCGB1D2, SCGB1D4*) are also upregulated following hormonal exposure (P4-diff) and downregulated after withdrawal (Fig.1e, Extended Data Fig.3a-c). In contrast, genes specific to the ciliated lineages (*FOXJ1, PIFO, TPPP3*) are upregulated at P4-diff and remain expressed after hormonal withdrawal (Fig.1e). Whilst the stromal compartment is the primary sensor of progesterone decline^16^, it is unclear whether the epithelial compartment also responds to hormonal withdrawal. Comparison of expression signatures of EOs between Horm Withdr 48h and P4-diff results in 41 upregulated and 63 downregulated genes, suggesting that the effect of hormonal withdrawal is indeed modest (Extended Data Fig.3d, Supplementary Tables). Genes encoding for receptors of peptide hormones (*SSTR1*) and retinoic acid (*RXRG*), as well as those involved in regeneration in other tissues (*MMP10*) are upregulated^21^. In contrast, markers of secretory cells and antagonists of the WNT signalling pathway (*SFRP1*, *NOTUM*) are downregulated upon hormonal withdrawal (Extended Data Fig.3d, Supplementary Tables).

Genes found at EOs at Post-breakdown 24h compared to the corresponding hormonally treated EOs that did not undergo breakdown (NB_H) are involved in response to wounding (Fig.1f, Supplementary Tables). There is a robust upregulation of cytokines associated with recruitment of immune cells (*CXCL1*, *CXCL6, CXCL8, CXCL10, IL23A*), genes involved in wound healing (*FGF2, AREG*), fibrinolysis (*PLAU, PLAUR*), and tissue remodelling (*MMP7, ADAMTS9, ITGA2*) (Fig.1e-g, Supplementary Tables). Many of these are characteristic of the menstrual window^22,23^. PLAUR, the plasminogen activator receptor that plays a key role in fibrinolysis, is localized to the shedding functional endometrial layer as well as in EOs specifically at Post-breakdown 24h (Fig.1h)^24^. In response to breakdown, there is also an increase in expression of *SOX9* and *FUT4* (coding for the surface antigen SSEA-1), potential markers of endometrial stem/progenitor cells^17^ (Extended Data Fig.4a-c). Novel genes found at this crucial time point are *THBS1* and *KRT13*, involved in wound healing in the intestine and lung respectively^25,26^ (Extended Data Fig.4a, d, e).

To further explore how hormonal treatment before breakdown affects endometrial regeneration, we compared EOs at Post-breakdown 24h to control EOs which had not been exposed to hormones before breakdown (B_NH) at the same timepoint (Extended Data Fig.5a). EOs exposed to hormones upregulated several acute phase inflammatory genes (*SAA1, SAA2, DUOX2*), and genes related to leukocyte migration (*IL10, CCR7, CCL20, TREM1, PF4V1, IRAK3*) at Post-breakdown 24h (Extended Data Fig.5a-b, Supplementary Tables). Overall, we observed better recovery of EOs undergoing the full IVMC protocol involving both hormonal treatment and mechanical breakdown (Extended Data Fig.5c), further confirming the physiological relevance of our model.

Re-exposure of EOs to estrogen for 48h (E2-diff 48h) attenuates the gene signature associated with response to breakdown and reactivates the expression of hormonally responsive genes (*PGR, GREB1, COL1A2*, *OLFM4*) and proliferative markers (Fig.1e, Extended Data Fig.3e-f, Supplementary Tables). The transcriptomic shift at this timepoint indicates the onset of the following cycle, reflecting the transition to the proliferative phase.

In summary, this IVMC protocol faithfully models the epithelial dynamics of repeated menstrual cycles, replicating key gene signatures and processes observed *in vivo*.

### Regenerating EOs after breakdown resemble the *in vivo* luminal epithelium of the menstrual and proliferative phases

In order to identify the key drivers in regeneration of the organoids, we next sought to investigate the *in vivo* epithelial cell states that are represented in EOs undergoing the IVMC protocol after mechanical breakdown. We first integrated epithelial cells from our single-cell RNA sequencing (scRNAseq) dataset of endometrial biopsies and whole uteri with other datasets (Wang *et al.*, Lai *et al.*, Huang et al. and Shih *et al.*)^13,27–30^ (Extended Data Fig.6a). These datasets together span the entire menstrual cycle including the menstrual phase (Extended Data Fig.6b). To align the IVMC with *in vivo* dynamics, we reanalysed the data and re-annotated the epithelial cells considering their temporal emergence across the menstrual cycle, enabling us to identify distinct epithelial populations specific to each phase. We identify 14 epithelial clusters, two ciliated populations and 12 non-ciliated (Fig.2a, b). The ciliated populations align with previously annotated clusters, including a pre-ciliated population enriched during the proliferative phase (*CCNO, CDC20B, MAD2L1, NEK2*), and a mature ciliated population (*PIFO, FOXJ1, RSPH1, TPPP3*)^13,31^. Among the non-ciliated populations there are hormone responsive (early and late), secretory (early, mid, late), cycling, luminal (early and late) cells and a few cells expressing *KRT5* and *MUC5B* (Fig.2c, Extended Data Fig.6d, Supplementary Tables)^13,31,32^. We also identify a transcriptionally active cluster characterized by high intronic fraction and genes involved in RNA splicing (*PNN*, *CCNL1, FUS*), and the Hippo signalling pathway (*TEAD1*, *WWC1*) (Fig.2c, Extended Data Fig.6c, Supplementary Tables). Lastly, we also identify a novel cluster annotated as ‘remodelling’ because it appears in the early phases of the menstrual cycle, has high vimentin (*VIM)* expression, but also expresses the epithelial marker, *EPCAM* (Fig.2c, Extended Data Fig.6e, f, Supplementary Tables). VIM positive cells can be identified *in vivo* specifically on the luminal surface in secretory phase and on the apical side of glands in menstrual phase (Extended Data Fig.6g). With this new analysis, we have generated a map of epithelial cell states across the different phases of the menstrual cycle.

**Figure 2:**
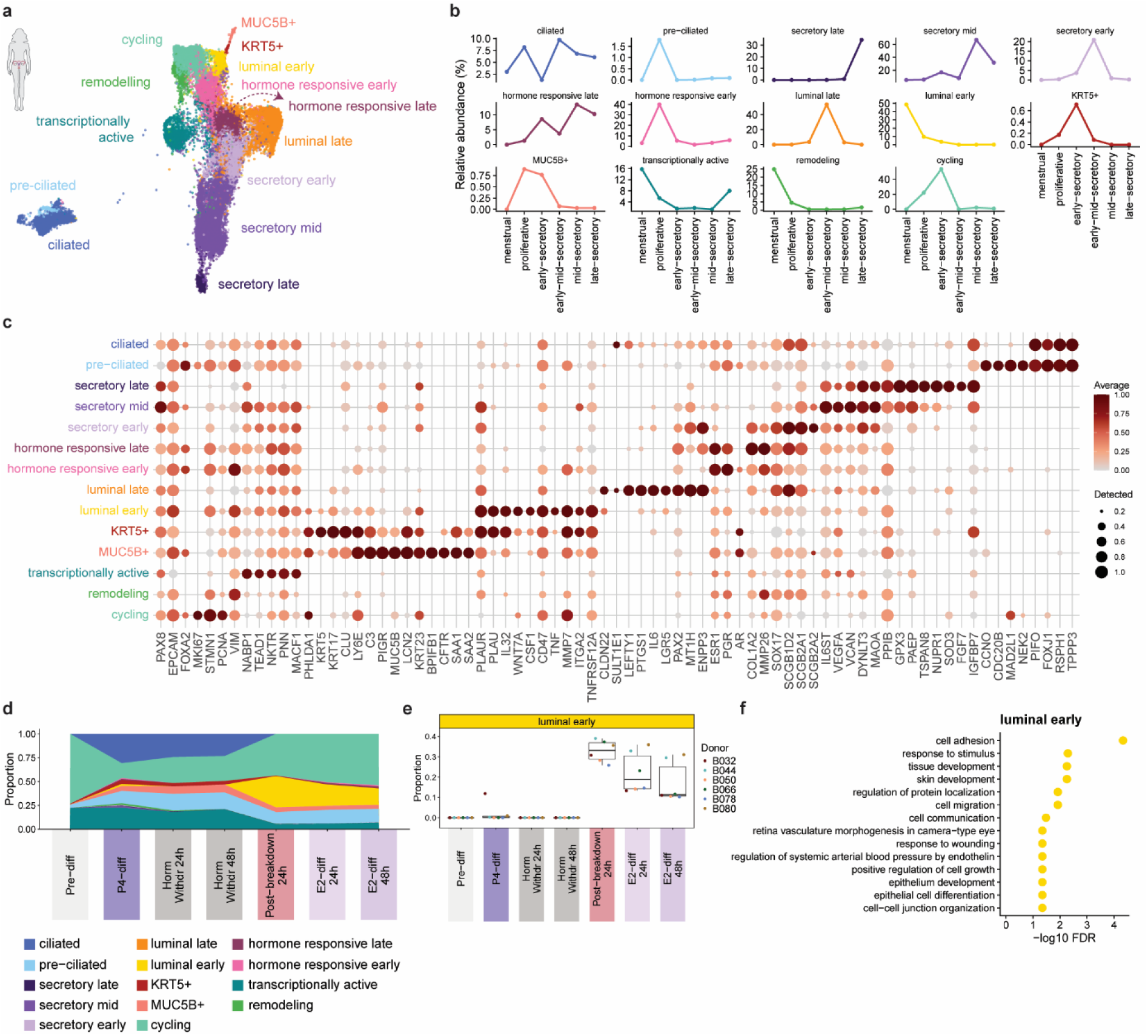
Regenerating EOs after breakdown resemble the luminal epithelium of the menstrual and proliferative phases. (**a**) UMAP visualization of epithelial cell clusters from integrated scRNAseq data of endometrial tissue across the menstrual cycle (n=23 donors). (**b**) Relative abundance plots of all *in vivo* epithelial cell clusters across phases of the menstrual cycle. (**c**) Dot plot illustrating the expression of selected marker genes in each *in vivo* epithelial cell cluster. Dot size represents the proportion of expressing cells, while colour denotes log2-transformed expression levels, normalised across all cell clusters. (**d**) Area plot showing the relative abundance of the deconvolved *in vivo* epithelial cell clusters represented in EOs across the IVMC protocol, averaged across all donors. (**e**) Box plot quantifying the estimated proportion of the *in vivo* luminal early cluster represented in EOs across the IVMC protocol. Individual dots represent individual donors (n=6 independent EO lines). (**f**) Biological processes enriched in the top 100 marker genes for the *in vivo* luminal early cluster.

To determine the cell states associated with epithelial regeneration after breakdown, we performed a deconvolution analysis of our *in vitro* bulk RNAseq data across the IVMC timepoints using this re-analysed *in vivo* scRNAseq dataset as a reference (Fig.2d). Cycling and luminal early clusters are the most abundant at the Post-breakdown 24h timepoint in all EO lines (Extended Data Fig.7). Whilst the proportion of cycling cells in EOs gradually increases in E2-diff 24h and E2-diff 48h phases (Fig.2d), the luminal early cluster is most abundant at the Post-breakdown 24h phase, representing ∼40% of the cells at this timepoint and then decreasing during E2-diff phases (Fig.2e). This luminal early cluster dominates during the menstrual phase *in vivo* (∼50% of cells), while its abundance drops drastically in the secretory phase (Fig.2b). It expresses many genes associated with response to wounding (*PLAU, MMP7, CSF1, ITGA2, CD47, WNT7A)* (Fig.2c, f, Supplementary Tables). *ITGA2,* involved in cell adhesion and migration and marking the luminal progenitor cells in the mammary gland, is highly expressed in the luminal early cell cluster ^33^. ITGA2 is not expressed by secretory phase luminal epithelium but is detected in both luminal and glandular cells during the menstrual phase (Extended Data Fig.6h). *CD47,* another marker of the luminal early cluster, is implicated in murine intestinal wound healing and is more strongly detected in the luminal epithelium during the menstrual than in the secretory phase (Extended Data Fig.6i)^34^. In contrast, the luminal late cluster is abundant during the mid-secretory phase (Fig.2b) and expresses receptivity associated genes (*IL6*, *LGR5*, *ENPP3, SCGB1D2)* in keeping with roles in embryo implantation (Fig.2c, Supplementary Tables). Indeed, we find that ENPP3 is present in the luminal epithelium in the secretory but not in the menstrual phase (Extended Data Fig.6j). In our data, *LGR5*, previously described as a marker of the luminal epithelium, is highly expressed in the luminal late cluster with transcripts being detected in the secretory but not menstrual phase (Fig.2c and Extended Data Fig.6k)^9,13,31^.

Taken together, these findings reveal that the luminal epithelium is dynamic, acquiring a wound healing profile in the menstrual phase and a receptivity associated profile in the secretory phase. In response to breakdown, EOs undergoing the IVMC protocol resemble the *in vivo* luminal early epithelium at a transcription level.

### *WNT7A*, a marker of the luminal early epithelium, is temporally associated with the response to breakdown *in vitro*

Our results thus point to a key role for the luminal early epithelium in the response to breakdown. We next sought to investigate what drives this response. Among the genes expressed in the luminal early cluster (Fig.2c), *WNT7A*, a ligand of the WNT signalling pathway, is a potential candidate for its role in epithelial regeneration (Fig.3a). *WNT7A* transcripts are present on the luminal epithelium in the menstrual phase, as previously reported^35^ (Fig.3b). Interrogation of WNT7A dynamics across the IVMC protocol reveals a transient peak in expression at Post-breakdown 24h after mechanical disruption compared to stably low expression in control EOs that did not undergo breakdown (NB_H) (Fig.3c). This pattern is consistently observed in EOs in continuous menstrual cycles (Fig.3d). Spatially, *WNT7A* transcripts are undetectable in EOs at Horm Withdr 48h but are uniformly present at Post-breakdown 24h (Fig.2e, f).

**Figure 3:**
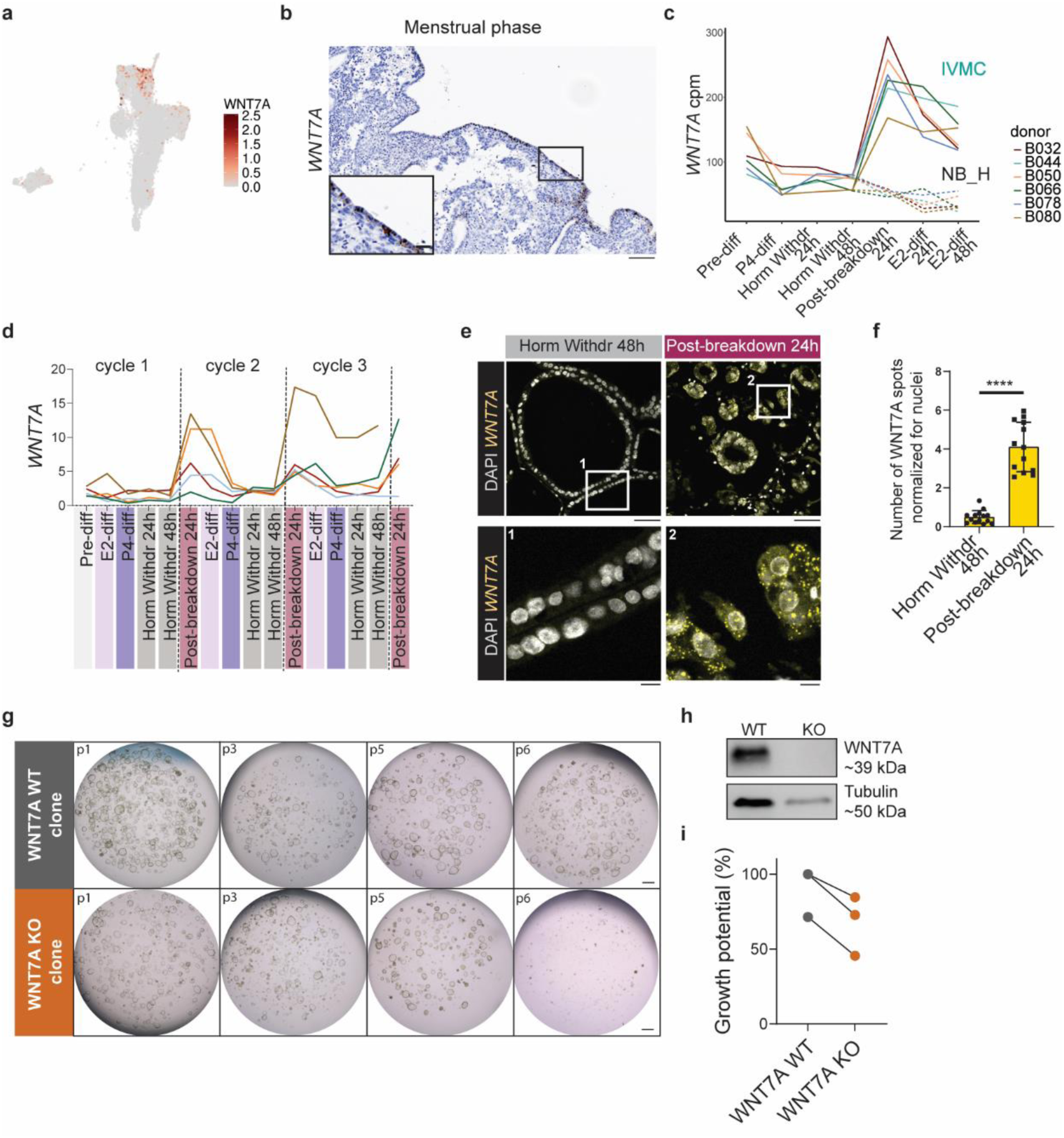
***WNT7A*, a marker of the luminal early epithelium, is temporally associated with the response to breakdown *in vitro*.** (**a**) UMAP visualization showing the log₂-transformed expression of *WNT7A* across individual cells *in vivo*. (**b**) *In situ* hybridization (ISH) showing localization of *WNT7A* transcripts in menstrual phase endometrium (n=1 donor). Scale bars, 100 μm (main), 50 μm (inset). (**c**) Line plots depicting cpm of *WNT7A* in batch corrected samples from bulk RNAseq data. Solid lines represent expression in EOs undergoing the IVMC protocol. Dashed lines represent expression in control EOs (NB_H) (**d**) qRT-PCR analysis of *WNT7A* expression of EOs undergoing continuous cycles of the IVMC protocol, normalised to housekeeping genes (n=5 independent EO lines). (**e**) ISH showing localization of *WNT7A* transcripts in EOs at Horm Withdr 48h and Post-breakdown 24h timepoints. Scale bars, 50 μm (main), 10 μm (inset). (**f**) Quantification of *WNT7A* ISH signal from (e). Significance was assessed using unpaired t-tests. Asterisks (****) correspond to *P < 0.0001*. (**g**) Representative brightfield images of WNT7A WT and WNT7A KO clones over six passages. Scale bar, 500 μm. (**h**) Western blot for WNT7A protein in WNT7A WT and WNT7A KO clones depicted in (g). (**i**) Percentage of WNT7A WT and WNT7A KO clones that could be frozen down as an estimate of growth potential (n=3 independent EO lines). Abbreviations: NB_H; No Breakdown Hormones.

The temporal and spatial regulation of *WNT7A* in EOs after breakdown suggests a role in epithelial renewal. In mice, WNT7A has an essential role in uterine glandular development^36^. In humans, *WNT7A* has been suggested to be implicated in both endometrial regeneration and the secretory transformation of the epithelium because it is downregulated by progesterone^35^, as also seen in our data (Fig.3c, d). To investigate whether WNT7A has a direct effect on the epithelium we generated WNT7A knockout (KO) organoids using CRISPR gene editing and assessed the impact on epithelial regeneration. Long-term survival of WNT7A KO clones is severely compromised compared to wild type (WT) clones (Fig.3g-h). Clonal cultures cannot be expanded beyond 5-7 passages after their establishment (Fig.3g). The overall growth potential, defined by the ability to expand and freeze down clones, is clearly lower in WNT7A KO clones (Fig.3i).

In summary, these findings confirm that *WNT7A*, a marker of luminal early cells in menstrual phase *in vivo*, is transiently induced in EOs in response to breakdown *in vitro*. Failure of EOs to survive long-term in the absence of WNT7A, suggests it has a role in endometrial renewal.

### Transient states of ciliated and non-ciliated cells emerge in EOs during the regenerative window, with *WNT7A* linking breakdown to proliferation

Limited accessibility of the endometrium during menstruation has hindered our understanding of how it rapidly regenerates in each cycle. We have now pinpointed *WNT7A*-expressing luminal cells as critical drivers for long-term epithelial maintenance. We next explored the emergence of distinct epithelial states and the identity of *WNT7A-*expressing cells in response to breakdown. We adjusted our *in vitro* protocol with finer time resolution after breakdown to recapitulate this crucial part of the cycle and performed scRNAseq of three independent EO lines (Extended Data Fig.8a-c). EOs were collected at 48 hours after hormonal withdrawal (Horm Withdr 48h), at 12, 24, 48 hours after breakdown (Post-breakdown 12h, Post-breakdown 24h, Post-breakdown 48h) and then at 24 and 48 hours after treatment with estrogen (E2-diff 24h, E2-diff 48h) (Fig.4a).

**Figure 4:**
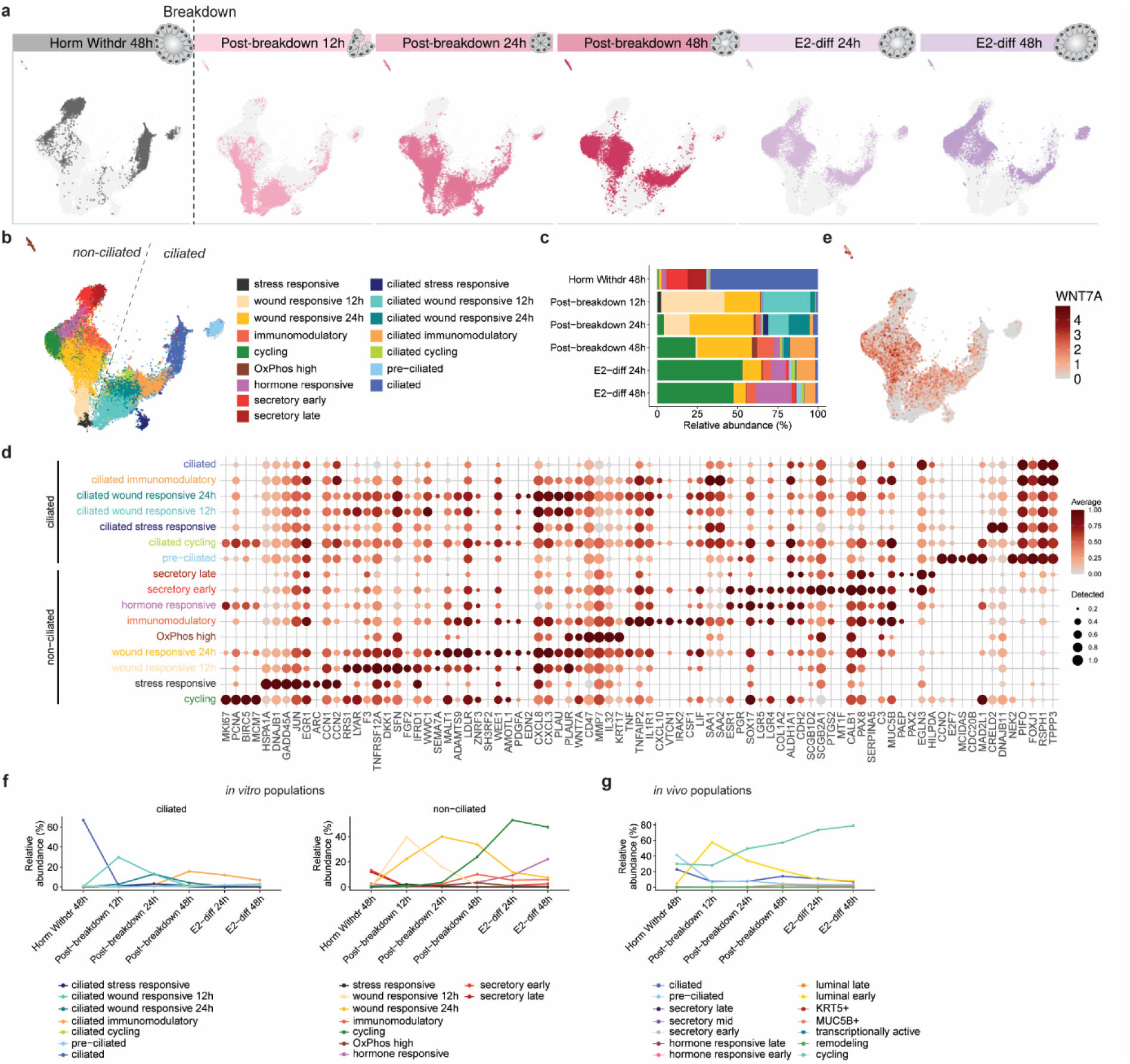
Transient states of ciliated and non-ciliated cells emerge in EOs during the regenerative window, with WNT7A linking breakdown to proliferation. (**a**) UMAP visualizations of cells coloured for each timepoint in separate panels. EOs are subjected to the IVMC protocol and collected for scRNAseq before breakdown (Horm Withdr 48h), after breakdown (Post-breakdown 12h, 24h, 48h) and after estrogen treatment (E2-diff 24h, E2-diff 48h) (n= 3 independent EO lines). (b) UMAP visualization of cells coloured by annotated clusters. (**c**) Bar plot showing the relative abundance (in percentage) of each cell cluster in the different timepoints. (**d**) Dot plot illustrating the expression of selected marker genes in each epithelial cell cluster. Dot size represents the proportion of expressing cells, while colour denotes log2-transformed expression levels, normalised across all cell populations. (**e**) UMAP visualization showing the log₂-transformed expression of *WNT7A* across individual cells. (**f**) Line plot showing the relative abundance (in percentage) of each *in vitro* cell cluster in the different timepoints. (**g**) Line plot showing the relative abundance (in percentage) of each predicted *in vivo* cell cluster corresponding to cells across the different timepoints *in vitro*.

We identify 16 distinct clusters segregating into seven ciliated and nine non-ciliated sub-clusters (Fig.4b, Extended Data Fig.8d, Supplementary Tables). Before breakdown (Horm Withdr 48h), differentiated ciliated and secretory cell populations are abundant (Fig.4c). There is a higher proportion of ciliated cells at this timepoint (Fig.4c), in agreement with our earlier observation that ciliated markers are not downregulated following hormonal withdrawal, unlike secretory markers (Fig.1e). In the immediate response to breakdown, we first identify non-ciliated stress responsive clusters appearing at Post-breakdown 12h with high expression of heat shock proteins (*HSPA1A, DNAJB1*) (Fig.4d, Extended Data Fig.8d-e, Supplementary Tables). Stress responsive ciliated cells appear later at Post-breakdown 24h and specifically express genes involved in the unfolded protein response (*CRELD2, DNAJB11*), indicative of a cellular adaptation to stress (Fig.4d, Extended Data Fig.8d, Supplementary Tables). In addition, we detect ciliated and non-ciliated cells, which we annotate as wound responsive 12h, due to their increased expression of injury response genes (*TNFRSF12A*, *SFN*), genes involved in remodelling (*F3*, *FGF2*) and in ribosome biogenesis and rRNA processing (*LYAR, RRS1*) (Fig.4d, Extended Data Fig.8d-e, Supplementary Tables). At Post-breakdown 24h, both ciliated and non-ciliated cells are annotated as wound responsive 24h. They acquire a different transcriptomic profile, expressing genes also identified in our bulk RNAseq analysis (*PLAUR, CXCL8, KRT17, MMP7*) (Fig.1e), involved in wound healing processes such as cell migration, adhesion and cytoskeleton organization (Fig.4d, Extended Data Fig.8d-e, Supplementary Tables).

At Post-breakdown 48h, a distinct cluster with high expression of genes involved in oxidative phosphorylation is present (Extended Data Fig.8d-e, Supplementary Tables). This cluster, which we annotate as OxPhos high, also expresses genes specific to the luminal early cluster *in vivo* (*WNT7A*, *IL32, MMP7, CD47*, and *KRT17)* (Fig.4d, Supplementary Tables). Additionally, clusters with immunomodulatory function are found at the same timepoint within both the ciliated and non-ciliated compartments, expressing genes of the TNF family (*TNF, TNFAIP2*), cytokines (*CXCL10, LIF*), cytokine receptors (*IL1R1*) and acute inflammatory response genes (*SAA1, SAA2*) (Fig.4d, Extended Data Fig.8d-e, Supplementary Tables). Cycling cells, found in S and G2M phases of the cell cycle, are only emerging from Post-breakdown 48h onwards (Extended Data Fig.8d-g). Most cycling cells are found in the non-ciliated (20%) and only sparsely in the ciliated (less than 2%) compartments (Extended Data Fig.8d). This suggests that non-ciliated cells may be the main contributor epithelial regeneration after breakdown.

Following re-exposure to E2, a hormone responsive population appears, marked by the expression of *ESR1* and *PGR*, alongside an early secretory cluster (*SCGB2A1, SCGB1D2*) (Fig.4d, Extended Data Fig.8d-e, Supplementary Tables). Due to the absence of additional hormonal stimuli (P4, PRL, cAMP), the *PAEP* expressing secretory late cluster remains scarce as expected. Within the ciliated compartment, E2 treatment increases the abundance of the pre-ciliated cell cluster (*FOXJ1, CCNO, MCIDAS, CDC20B*) (Fig.4d, Extended Data Fig.8d, Supplementary Tables). This cluster also shows unique enrichment for *E2F7*, a transcriptional regulator of canonical S-to-G2 progression that promotes multiciliated cell differentiation^37^ (Fig.4d).

Our data reveals that *WNT7A* is highly expressed in both ciliated and non-ciliated wound responsive 24h and cycling clusters, while it is only lowly expressed in wound responsive 12h clusters and completely absent from the stress responsive clusters (Fig.4d, e). This further confirms the temporal regulation of *WNT7A in vitro*. We show that in response to breakdown, EOs gradually go through states of immediate stress response followed by wound healing, with *WNT7A* expression peaking at 24h, before the onset of cell proliferation (Fig.4f). Interrogation of the *in vivo* epithelial identities in EOs undergoing the IVMC protocol further suggests that the luminal early cluster, expressing *WNT7A*, is similarly enriched before cycling cells become dominant (Fig.4g).

Overall, our analysis underscores the ability of our IVMC protocol to capture transient states emerging in response to breakdown. Both ciliated and non-ciliated cells initially adopt stress-responsive states, followed by a shift toward wound-responsive programs characterised by induction of cytokines and genes involved in cell adhesion and migration followed by proliferation. *WNT7A*, is widely expressed by both ciliated and non-ciliated cells before the onset and during proliferation further reinforcing the significance of the luminal epithelium in coordinating endometrial regeneration.

### The luminal early epithelium is a key signalling mediator during endometrial regeneration

Our findings highlight the importance of the luminal epithelium expressing *WNT7A* during mucosal regeneration. As menstruation and regeneration are complex processes involving many cell types, we investigated how the luminal epithelium may orchestrate this process through its interactions with other endometrial cells. To achieve this, we performed CellChat analysis of all cells in our integrated scRNAseq *in vivo* endometrial dataset. We specifically selected cells from the menstrual and early proliferative phases of the cycle to focus on the regeneration window. Defining the luminal early population as the ‘sender’, we confirm that it is predicted not only to interact with other epithelial populations but also with immune, mesenchymal and endothelial cells (Fig.5a). The highest interaction probability is between luminal early epithelial cells and venous endothelium (Fig.5a).

Multiple signalling networks including CXCL, GDF, VEGFA, CALCR, LIFR, Virsfatin, SEMA3 and TNF may mediate the interaction between luminal early epithelial cells and endothelial cells (Fig.5b). We focussed on CXCL interactions as its ligands are upregulated in EOs at Post-breakdown 24h (Fig.1e, Extended Data Fig.9a). *CXCL8* has a role in neutrophil recruitment and promotion of angiogenesis during the perimenstrual window^23,38^. It is expressed by luminal early cells *in vivo* and peaks at Post-breakdown 24h *in vitro* (Fig.5c-d, Extended Data Fig.9a). We confirm that EOs secrete high concentrations of IL-8 (encoded by *CXCL8*) at Post-breakdown 24h with EOs from the IVMC secreting higher amount compared to control EOs not treated with hormones (B_NH) (Fig.5e). This aligns with the enhanced inflammatory-like response, characteristic of the perimenstrual phase *in vivo*, which we detect in EOs subjected to the IVMC protocol at Post-breakdown 24h (Extended Data Fig.5a-b)^39^. The action of IL-8 can be mediated through its cognate receptors CXCR1, CXCR2 or the atypical chemokine receptor 1 (ACKR1)^38^. *ACKR1* is specifically expressed by venous endothelial cells (Fig.5c). Detection of CXCR1 positive epithelial cells in menstrual phase *in vivo*, as well as *in vitro* suggests that IL-8 might also directly affect the epithelium (Extended Data Fig.9b).

Recruitment of macrophages and neutrophils is characteristic of menstrual phase endometrium^39^. We identify interactions between luminal early cells and myeloid cells, mediated through Annexin, CSF, Complement, TNF and MIF signalling networks (Fig.5b, Extended Data Fig.9c). Annexin1 (ANXA1) is essential for the resolution of inflammation^40^. *ANXA1* is highly expressed in luminal early cells *in vivo* and upregulated at Post-breakdown 24h *in vitro* (Fig.5c-d). EOs subjected to the IVMC protocol have more ANXA1 positive cells compared to control organoids that have been broken down but not treated with hormones (B_NH) (Fig.5f).

Interactions between the luminal early and other epithelial cells involve the IL-6, WNT, PTN, MK and EGF signalling networks (Fig.5b, Extended Data Fig.9c). *WNT7A* is the main ligand of the WNT signalling pathway secreted from the luminal early epithelium as well as from EOs, at the Post-breakdown 24h timepoint (Fig. 5c, Extended Data Fig.10a). WNT7A can acts through various frizzled (FZD) receptors^41–43^. Among them, *FZD6* is highly expressed in various epithelial cell types and both venous and arterial endothelial cells *in vivo* (Fig.5c). EOs express higher levels of *FZD6* and *FDZ10* at Post-breakdown 24h (Extended Data Fig.10b-d). While *FZD6* is widely expressed in EOs upon breakdown, *FZD10* is mainly expressed from non-ciliated cells (Extended Data Fig.10e). How WNT7A-responsive endometrial cells coordinate the regenerative process at menstruation will need further investigation.

**Figure 5:**
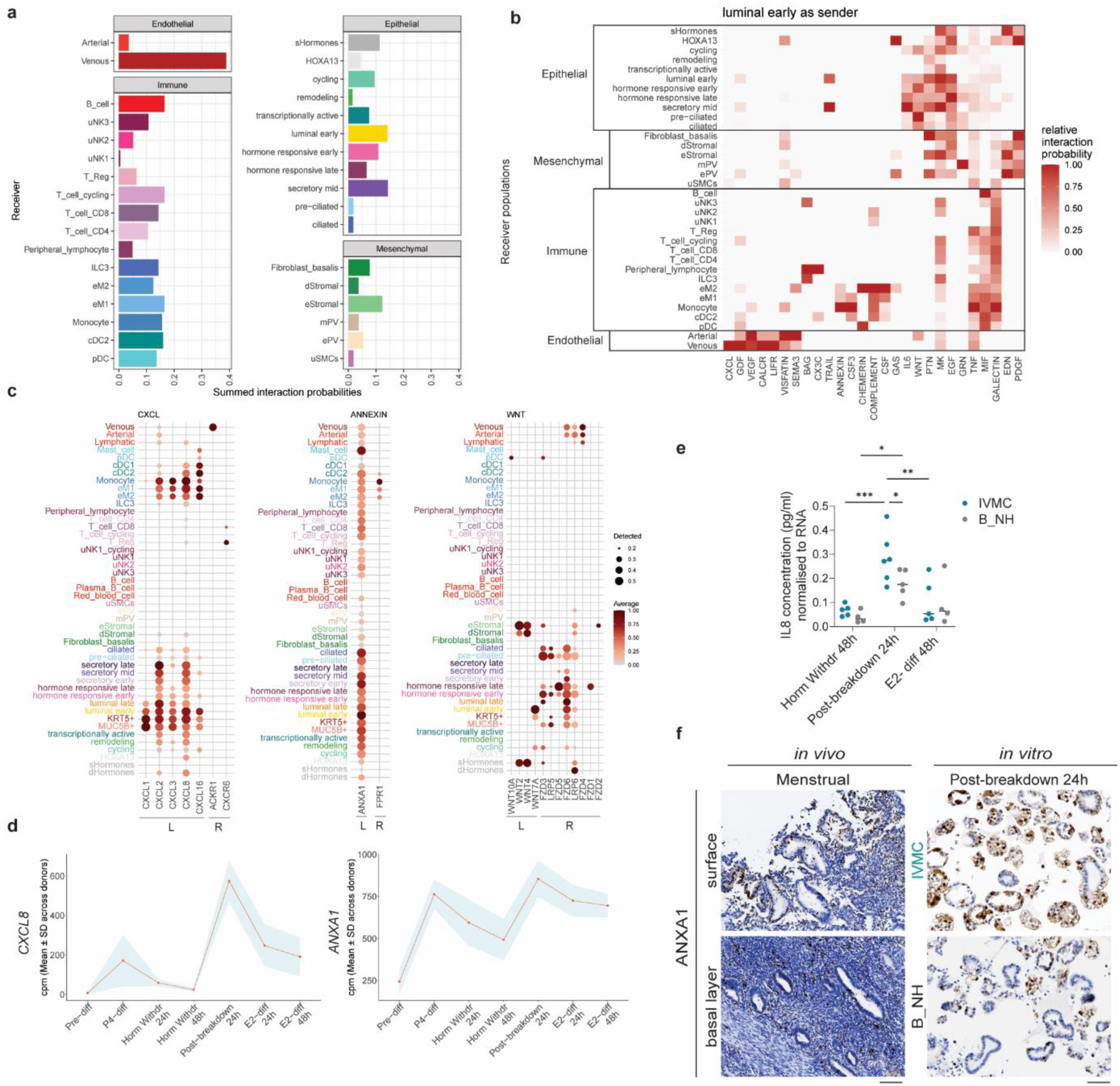
The luminal early epithelium is a key signalling mediator during endometrial regeneration. (**a**) Total estimated interaction probabilities between luminal early epithelial cells (sender) and all the other epithelial and non-epithelial cell clusters (receiver). (**b**) Heatmap showing the estimated relative interaction probabilities for different signalling networks between luminal early epithelial cells (sender) and all the other epithelial and non-epithelial cell clusters (receiver). (**c**) Dot plot illustrating the expression of ligands (L) and receptors (R) involved in CXCL, Annexin and WNT signalling networks. Dot size represents the proportion of expressing cells, while colour denotes log2-transformed expression levels, normalised across all cell populations. (**d**) Line plots showing the mean expression levels in cpm of *CXCL8* and *ANXA1* across the IVMC protocol. The red line represents the mean cpm values across donors (n=6 independent EO lines), while the shaded blue region indicates the standard deviation around the mean. (**e**) Concentration (pg/ml) of IL8 in supernatants of EOs in the IVMC protocol and control EOs (B_NH) at different timepoints. Concentration is normalised to RNA concentration per well to account for organoid confluency. Statistical analysis was performed using two-way ANOVA (hormonal treatment × timepoint), with Tukey’s multiple comparisons test. Asterisks indicate significance level *P* < 0.05 (*), *P* < 0.01 (**), *P* < 0.001 (***). (**f**) Representative IHC image for ANXA1 staining in sections from functional and basal layers of menstrual endometrium (n=4 donors for each phase) and EOs at Post-breakdown 24h (n=4 independent EO lines). Scale bars of tissue sections, 100 μm. Scale bars of EO sections, 50 μm. Abbreviations: B_NH; Breakdown No Hormones.

Taken together, these findings define the pivotal role of the luminal epithelium in coordinating the response to breakdown and regeneration through multiple signalling pathways to both epithelial and non-epithelial endometrial cells.

## Discussion

In this study, we have used EOs to establish an *in vitro* menstrual cycle (IVMC) protocol that faithfully and repeatedly recapitulates the sequential epithelial phases of hormonal differentiation, breakdown and regeneration. This model, together with the *in vivo* reference of epithelial cell states across the cycle, has allowed an in-depth study of the critical period spanning menstrual shedding and epithelial renewal. We uncover the cellular dynamics and transcriptional transitions in response to breakdown that underpin endometrial epithelial regeneration.

Beyond modelling cyclical dynamics, the IVMC provides a powerful platform to investigate how distinct epithelial populations emerge during the menstrual cycle. Prior to hormonal differentiation, EOs are proliferative and express *CDH2* and *ALDH1A1,* markers of glandular epithelium in the basal layer that is not shed and which is thought to be the primary source of epithelial regeneration following menstruation^18,19^. Using the IVMC protocol, we aimed to better define the regenerative cell states after breakdown. While significant work has been made by us and others to define the distinct epithelial cell states in the endometrium using transcriptomic technologies, the usage of different markers to define these states poses challenges for harmonizing and integrating findings^13,27,31^. Here, we defined epithelial subtypes based on their emergence during the menstrual cycle, enabling us to better correlate *in vitro* with *in vivo* cell states. Using full thickness uterine FFPE tissue blocks carefully dated to span the menstrual window, we validate the key cell states and their localization *in vivo*. Our data revealed a clear enrichment of the luminal early epithelial signature (*WNT7A, ITGA2, CD47*) in EOs after breakdown, supporting an understudied idea that luminal epithelial cells could also participate in regeneration^6,7^. This is further supported by scanning electron microscopy studies showing re-epithelialization from the unshed surface epithelium^7^. Furthermore, separation of functional from basal endometrial layers by laser capture microdissection followed by transcriptomic analysis also found that shedding functional zones express genes involved in tissue degradation and regeneration^6^, as we find in EOs after breakdown. Lineage-tracing experiments using the IVMC protocol will be essential to determine the long-term fate of individual cell populations, the relative contributions of basal versus luminal compartments and the capacity of committed cell types to contribute to epithelial regeneration, potentially via dedifferentiation as seen in the intestine and lung ^44,45^.

*WNT7A* emerges as a defining marker of genes upregulated in the regenerative window in IVMC. *WNT7A* expression is tightly linked to menstrual and proliferative phase luminal epithelium *in vivo* and temporal profiling *in vitro* shows upregulation after breakdown. While prior studies in mice have implicated *WNT7A* in uterine patterning^46^, postnatal gland formation^36^ and anti-apoptotic function^47^, its role in the human endometrium remained undefined. In other adult tissues, WNT7A plays roles in epithelial integrity, maintenance of stem cells, wound healing and inhibition of inflammatory stimuli through diverse mechanisms ^41,42,48–52^. Here, we show that *WNT7A* is a potential regulator of endometrial epithelial regeneration as its genetic deletion in EOs impaired long-term survival. Our findings position WNT7A as a central epithelial-derived cue orchestrating survival and repair following menstrual breakdown, providing a foundation for future work to dissect its downstream effectors and responsive cell populations.

Cell–cell communication analysis of the human endometrium during the menstrual and proliferative phases predicts that luminal epithelial cells function as a key signalling hub during regeneration. This complements previous findings demonstrating interactions between cells of non-epithelial origin in endometrial regeneration^31^. Our analysis revealed significant predicted interactions between luminal early epithelial and endothelial cells, particularly mediated by cytokines of the CXCL family such us CXCL8 (IL-8) that functions in recruitment of neutrophils and angiogenesis during menstruation^23^. IL-8 secretion by EOs following breakdown in the IVMC model further supports the physiological relevance of our system. CXCR1, the IL-8 receptor, is found on EOs post-breakdown, suggesting that IL-8 could act in an autocrine or paracrine manner on the epithelium itself. Our analysis also suggests interactions between epithelial and stromal cells, although these have not been functionally validated. Co-culture systems would greatly enhance our understanding of all these^53^.

In summary, our IVMC protocol establishes a physiologically relevant, tractable system to interrogate the epithelial cell states across the menstrual cycle that are otherwise challenging to capture. By recapitulating the cyclical dynamics of the menstrual cycle *in vitro*, our organoid model not only illuminates the phases of tissue regeneration in response to breakdown but also provides a powerful platform for dissecting the molecular and cellular mechanisms of this remarkable process of scarless repair. Furthermore, the IVMC approach, in combination with existing *in vivo* data, will enable more accurate modelling of how the menstrual cycle dynamics are disrupted in gynaecological disorders including endometriosis^28,32,54,55^, Asherman syndrome^56^ and polycystic ovary syndrome (PCOS)^57^. These investigations will allow understanding of disease mechanisms and possible therapeutic interventions.

## Methods

### Origin of samples

Human endometrial tissue blocks were kindly donated by Professor Ashley Moffett (University of Cambridge, UK). They were collected at Addenbrookes Hospital, Cambridge before 1 September 2006 (prior to the implementation of the Human Tissue Act) and therefore formal consent for research use was not required. These samples are categorized as ‘Existing Holdings’ under the Human Tissue Act and were therefore eligible for use in this project. The hysterectomy uteri span all phases of the menstrual cycle (dated by a pathologist) and are properly oriented to allow evaluation the full thickness of the endometrium.

Endometrial organoids were derived from biopsies of women in the secretory phase of the cycle undergoing IVF at the Bourn Hall Fertility Clinic with approval from East of England Cambridge Central Research Ethics Committee (17/EE/01551 IRAS 225205). The transfer and use of all the endometrial tissue samples together with the established endometrial organoid cultures for this study are authorized under a Material Transfer Agreements (MTA) between the University of Cambridge and the Friedrich Miescher Institute for Biomedical Research (Agreements No. G119629 and G108313).

### Expansion and differentiation of EOs

EOs were grown as previously described^12^. Briefly, EOs were grown in Matrigel (Corning, 356231) droplets supplemented with endometrial organoid medium (EOM). Components of EOM are available in Supplementary Table 1. Differentiation of EOs was achieved via hormonal stimulation. First, EOs were primed with 0.5 nM E2 for two days followed by four additional days of combination of 0.5 nM E2, 80 nM P4, 100 μg/ml cAMP and 20 ng/ml PRL (Supplementary Table 1). For the IVMC protocol, hormones were withdrawn the following two days with EOM being changed every 24 hours. After hormonal withdrawal, EOs were broken mechanically with automatic (300 times) and manual (100 times) pipetting. EOs were then left to grow for 24 hours (for bulk RNA sequencing) or 48 hours (single cell RNA sequencing) in the absence of hormones. Another round of E2 treatment was then re-initiated for two days.

### Single cell dissociation

EOs were collected and Matrigel was removed using Cell Recovery Solution (Corning, 354253) for 50 min maximum on ice. EOs were washed with cold PBS pH (7.4) (Gibco, 10010-015) and broken apart by automatic pipetting (400 times). EOs were, first incubated with pre-warmed accutase cell detachment solution (Gibco, A11105-01) for 5 minutes at 37°C and next incubated with collagenase V (Sigma-Aldrich, C-9263) diluted in 10% FBS/Advanced DMEM/F12 for 20 minutes at 37°C. The digest was passed through a 40 μm nylon mesh cell strainer to ensure single cell suspension, and the flow through was collected in Falcon round bottom tubes (Corning, 352235). If undigested organoid fragments were still present, the collagenase step was repeated. Cell pellets were diluted in EOM, and cells were counted using the NucleoCounter® NC-202™ Automated Cell Counter (Chemometec, NC-200). EOs growing from single cells were supplemented with 10µM Y-27632 (Supplementary Table 1) until the formation of cystic structures.

### Fixation and paraffin embedding of EOs

EOs were collected at different timepoints through the IVMC. Matrigel was removed using Cell Recovery Solution (Corning, 354253) for maximum 40 min on ice. EOs were washed with cold PBS pH (7.4) (Gibco, 10010-015) and fixed in 10% neutral buffered formalin solution (Sigma-Aldrich, HT5011) for 1h at RT. Fixed EOs were centrifuged at 0.3 rcf, washed twice in PBS and embedded in pre-warmed, liquid Histogel (Epredia, HG-4000-012). After solidification at RT, the Histogel dome containing EOs was transferred into a labelled histology cassette (Cell Path, EAM-0309-72B) and stored in 70% ethanol until dehydration and paraffin embedding. EOs were loaded in the retort of the HistoCore PEARL tissue processor (Leica, 14 0493 50667) using overnight standard tissue dehydration method until paraffin inclusion. The samples were then removed from the tissue processor, transferred in molten paraffin and attached to the cassettes using the HistoCore Arcadia Embedding Center (Leica, 14 0393 57262 and 14 0393 57257). After solidification of the paraffin blocks, microtome (Leica, 149MULTI0C1) sectioning was performed.

### Immunohistochemistry

Immunohistochemistry was performed on 3-μm-thick paraffin sections of both endometrial tissues and organoids using the Leica Bond RX automated stainer. As part of the automated staining, the sections were first deparaffinized with the BOND Dewax Solution (Biosystems Switzerland AG, AR9222) and cleared with 100% ethanol. Next, heat induced epitope retrieval was performed with BOND Epitope Retrieval Solution 1, pH6 (Biosystems Switzerland AG, AR9961) or BOND Epitope Retrieval Solution 2, pH9 (Biosystems Switzerland AG, AR9640) at 100°C for 10 to 20min depending on the primary antibody. Sections were incubated with appropriate primary antibody dilutions for 60 minutes at room temperature followed by washes with BOND Wash Solution 1X (BOND Wash Solution 10X Concentrate, Biosystems Switzerland AG, AR9590). Details of all primary antibodies are listed in (Supplementary Table 2). The binding sites of the primary antibodies were then revealed using the ready-to-use Bond Polymer Refine detection kit (Biosystems Switzerland AG, DS9800) containing peroxide block solution, post primary, polymer reagent, DAB chromogen and hematoxylin counterstain. Sections were then, dehydrated with consecutive 5-minute incubations in 70% ethanol, 95% ethanol and 100% ethanol. Last, sections were dried at room temperature prior to addition of Tissue-Tek® Glas™ Mounting Medium (Sakura, 1408N). Coverslips 24×50mm #1,5 (Epredia, BB02400500SC13MNZ0) were applied using Tissue-Tek® Glas™ g2-E2 instrument (Sakura). Stained sections were imaged with the Zeiss Axioscan Z1 slide scanner.

### *In situ* hybridization assays

*In situ* hybridization assays were performed on 5-μm-thick paraffin sections of endometrial tissues using the RNA scope high-definition brown assay (ACD, 322310) and sections of EOs using the RNA scope Multiplex Fluorescent V2 Assay (ACD, 323100), following the manufacturer’s instructions. For both assays, sections were first baked at 60 °C for 1 hour and dewaxed with xylene (Sigma-Aldrich, 534056), cleared in 100% ethanol and air-dried. The sections were then treated according to the standard protocol: 10 minutes in Hydrogen Peroxide, 15 minutes (tissue) or 10 minutes (organoids) in Target Retrieval reagent and 30 min (tissue) or 10 minutes (organoids) at 40°C in Protease Plus. Sections were then incubated with gene specific probes for 2 hours at 40 °C (Supplementary Table 3) and signal was amplified with appropriate kit reagents. After signal amplification, tissue and organoid sections were processed differently according to the assay requirements. Tissue sections were treated with DAB for 10 min for signal visualization. Sections were then dehydrated, mounted in histological mounting media Histomount (HS-103, National Diagnostics) and imaged using the EVOS™ M5000 Imaging System (Thermo Fisher, AMF5000). After signal amplification, organoid sections were first fluorescently labelled for each probe using fluorophores; TSA Vivid 520 (ACD, 323271), TSA Vivid 650 (ACD, 323273) or TSA Plus Cyanine 3 (Akoya Biosciences, NEL744001KT) conjugated to the appropriate channel-specific amplifiers (C1/C2/C3). Sections were counterstained with DAPI (10 sec) and mounted with ProLong™ Gold Antifade Mountant (Life Technologies, P36934). Fluorescent signals were imaged using Stellaris 5 confocal microscope.

### Quantification of *WNT7A* transcripts

*WNT7A* transcripts from *in situ* hybridization assay were quantified in ImageJ (version 1.54f) by performing spot detection using the TrackMate plugin. DoG detector parameters were set to 0.852 micron for estimated object diameter and 1000 for the quality threshold. Simple LAP tracker was applied before spot quantification. Nuclei were segmented using the StarDist 2D plugin following default settings to be used for normalization. The number of spots was then normalized to the number of nuclei. In total, 14 image scans were used from EOs at Hormonal Withdrawal 48h and Post-breakdown 24h respectively. Significance values were determined using an unpaired t-test to compare the two groups.

### Illumina stranded mRNA-seq library preparation, bulk RNA-sequencing and analysis

cDNA libraries were prepared using the Illumina Stranded mRNA-Seq (Illumina IDT DNA/RNA UDI) preparation kit according to the manufacturer’s instructions. Libraries were pooled and sequenced with the Illumina NovaSeq 6000 platform, producing paired end reads of 50 base pairs each (2×50bp). Demultiplexing and FASTQ file generation were performed using Illumina’s bcl2fastq2 software. RNA-seq data was quantified on the transcript level using Salmon version (1.6.0)^58^ with parameters --gcBias --seqBias –numGibbsSamples=50. The reference index was constructed based on the transcriptome (transcripts.fa) from GENCODE release 38^59^, using the genome sequence as a decoy^60^ and a k-mer length of 23. Transcript-level estimated counts from Salmon were imported into R and summarized to the gene level using tximeta version (1.16.1)^61^. Raw count data were filtered to remove lowly expressed genes setting the following parameters: mincpm <-1 and minsamples <-x, where x is the number of different donors (endometrial organoid lines) of each dataset that was analysed. The filtered dataset was normalized using the TMM (Trimmed Mean of M-values) method with the calcNormFactors function in edgeR version (4.6.2)^62^ to account for differences in library size and composition. A generalized linear model (GLM) framework was used to model gene expression while accounting for donor effects. A design matrix was constructed as follows: *design ∼ donor + group* where "donor" accounts for individual variability and "group" represents the experimental conditions. DEA was performed using the quasi-likelihood F-test (QLF test). The glmQLFit function was used to fit the model, and the glmQLFTest function was applied with a contrast matrix to compare experimental conditions. Genes were classified as significantly upregulated or downregulated based on the following threshold: FDR < 0.05 and |logFC|>2 unless stated otherwise in figure legends. Data visualizations and all plots were generated using R version 4.5.0.

### Public scRNA-seq data

Processed data from the Reproductive Cell Atlas was downloaded from https://www.reproductivecellatlas.org/ and converted from h5ad to SingleCellExperiment format using the zellkonverter R package (v1.12.0)^63^. Cell-level annotations were extracted from the SingleCellExperiment object and incorporated into the analysis by matching cell barcodes. Cell cycle marker genes were obtained from https://github.com/ventolab/HECA-Human-Endometrial-Cell-Atlas/tree/main/utils.

### scRNA-seq reference index generation

The human genome sequence (GRCh38.primary_assembly.genome.fa) and transcript annotation (gencode.v38.annotation.gtf) files were downloaded from GENCODE release 38 ^64^. Coordinates for transcripts and introns were extracted from the gtf file using the getFeatureRanges function from the eisaR R package (v1.6.0)^65^, with arguments featureType = c("spliced", "intron"), intronType = "separate", flankLength = 50, 85 or 145 (depending on the read length), joinOverlappingIntrons = TRUE, collapseIntronsByGene = TRUE, keepIntronInFeature = TRUE. Sequences for the extracted features were obtained using the extractTranscriptSeqs function from the GenomicFeatures package (v1.46.1)^66^, and a splici index^67^ was generated using Salmon (v1.6.0)^58^ with the k-mer length set to 23 or 31, again depending on the read length. A feature-to-gene map was created using the getTx2Gene function from the eisaR package, adding an extra column indicating the feature type (spliced or unspliced).

### scRNA-seq processing (*in vivo* atlas)

Public scRNA-seq data were quantified using Salmon (v1.6.0) and alevin-fry (v0.4.3)^67^, in USA (unspliced-spliced-ambiguous) mode. Alevin-fry was run with parameters ‘-d fw --knee-distancè (for the generate-permit-list subcommand) and ‘--resolution cr-like --use-mtx‘ (for the quant subcommand). Initial quality control was performed using the alevinQC R package (v1.18.1)^68^.

Alevin-fry counts were imported into R (v4.4.2) using the loadFry function from the fishpond package (v2.12.0)^69^. Spliced and ambiguous counts were added up to form the main count matrix for downstream analysis.

To select only epithelial cells for further analysis, the raw counts were normalized using size factors calculated using the computeLibraryFactors function from the scuttle R package (v1.16.0)^70^ and the multiBatchNorm function from the batchelor R package (v1.22.0)^71^, using t he donor as the batch variable. The top 3,000 highly variable genes were defined using the modelGeneVar and getTopHVGs functions from the scran R package (v1.34.0)^72^, blocking on the data set in the variance calculation, and batch correction was performed using the fastMNN implementation from batchelor, again using the donor as the batch variable. Uniform Manifold Approximation and Projection (UMAP)^73^ was applied to the fastMNN reduced dimension output using the runUMAP function from the scater R package (v1.34.0)^70^, and a low-resolution clustering was applied to the UMAP representation using the makeSNNgraph function from the bluster R package (v1.16.0)^74^ and the cluster_louvain function from the igraph R package (v2.1.4)^75^ with the resolution parameter set to 0.001. The cluster with the largest over-representation of cells from the epithelial lineage, as determined by the Human Endometrial Cell Atlas (HECA) annotations, was retained for further analysis (n=85,354 cells). After subsetting to epithelial cells, only genes annotated as one of protein_coding (excluding ribosomal proteins, defined by the GO term "structural constituent of ribosome" (GO:0003735)), IG_V_gene, IG_C_gene, IG_J_gene, TR_C_gene, TR_J_gene, TR_V_gene, TR_D_gene, IG_D_gene, Mt_tRNA, Mt_rRNA were retained, leaving 20,231 of the original 60,649 genes. Quality metrics for cells, including the total number of UMI counts, the number of detected genes, and the fraction of the total UMI counts coming from mitochondrial genes, were calculated using the addPerCellQC function from scuttle. Further, the intronic fraction (unspliced UMI count / total UMI count) was calculated for each cell. Cells with total UMI count above 1,000, more than 500 detected genes, and mitochondrial count fraction below 25% were retained. In addition, samples with less than 30 cells remaining after these filtering steps were excluded, leaving 49,504 cells from 26 samples for further analyses.

Size factors were calculated using the computeSumFactors function from scran followed by multiBatchNorm from batchelor, using the donor as the batch annotation. 582 cells with size factors below 0.1 were excluded to reduce normalization-induced artifacts. Cell cycle assignment was performed by Seurat (v5.2.1)^76^, using the marker genes from the HECA atlas^31^. Next, we used geometric sketching to subsample cells from each donor, using the interface to the geosketch algorithm^77^ provided by the geosketch function from the sketchR R package (v1.2.0)^78^. For each donor, we extracted 2,000 highly variable genes using the modelGeneVar function from scran, which were used to perform principal component analysis using the runPCA function from scater, based on which the geosketch function was applied to select 10% of the cells (or 150 cells, whichever of the two numbers was largest). If the donor contributed less than 150 cells, all of them were kept. Based on the sketched cells (n=5,700), a new set of 3,000 highly variable genes were defined using modelGeneVar. These genes were used to define principal components using the multiBatchPCA function from batchelor, onto which all cells were projected before batch correction was performed using the reducedMNN function from batchelor. The resulting reduced dimension representation was used as the basis for t-SNE^79^ and UMAP projection, performed using the fitsne function from the snifter R package (v1.16.0)^80^ and the umap function from the uwot R package (v0.2.2)^81^, respectively. In both cases, the sketched cells were used to define the mapping to the low-dimensional space, which was then applied to the full set of cells via the project (snifter) and umap_transform (uwot) functions, respectively.

Clustering was performed on the sketched cells, using the clusterRows function from bluster, using a shared nearest neighbor graph (k=10, type="rank") and Leiden clustering^82^ with resolution 0.4. Cluster labels for the remaining cells were defined by assigning the cluster label of the nearest neighbor among the sketched cells. Clusters were manually annotated into 14 final classes. Finally, marker genes were calculated by comparing each pair of clusters using the pairwiseTTests function from scran, blocking on the sample and only considering upregulated genes, followed by calling combineMarkers with pval.type="all" to find specific marker genes. Only genes expressed in at least 0.05% of the cells were considered for the testing. Functional analysis of the top 100 marker genes for each cluster was performed using the gost function from the gprofiler2 R package (v0.2.3)^83^, using all tested genes as the universe and considering all gene sets from the GO:BP collection.

The processed *in vivo* epithelial atlas was used to deconvolve the timecourse bulk RNA-seq data, using the BayesPrism package (v2.2.2)^84^. Only protein-coding genes detected in at least 3 cells were considered, and marker genes were extracted using the BayesPrism functions get.exp.stat and select.marker with default settings.

### scRNA-seq processing (*in vitro* organoid time course)

Time course scRNA-seq data generated for this study were quantified using Salmon (v1.9.0) and alevin-fry (v0.8.0)^58^, in USA (unspliced-spliced-ambiguous) mode. Alevin-fry was run with parameters ‘-d fw --knee-distancè (for the generate-permit-list subcommand) and ‘--resolution cr-like-em --use-mtx‘ (for the quant subcommand). Initial quality control was performed using the alevinQC R package (v1.18.1)^59^. Following quantification, the scRNA-seq data was processed following the same steps as the in vivo data described above, with a few differences. First, the initial selection of cells from the epithelial lineage was skipped, and all cells were retained for the main analysis. Second, no batch correction was performed, effectively excluding the steps above corresponding to the multiBatchNorm (replaced by logNormCounts), multiBatchPCA (replaced by calculatePCA) and reducedMNN functions (all cells were annotated to the same batch). Leiden clustering was performed with resolution 0.125 and clusters were manually annotated into 16 final classes. In addition to the Seurat cell cycle annotation, tricycle was applied to obtain a continuous cell cycle position^85^. Sketching was applied to the full data set, extracting 10% of the cells (n=8,562).

Next, the SingleR R package (v2.8.0)^86^ was used to annotate the organoid time course data using the cell type labels defined in the in vivo atlas. The aggregateAcrossCells function was used to build a set of reference profiles from the in vivo data, followed by application of the SingleR function to annotate the cells from the organoid time course.

### Cell-cell communication analysis

In order to investigate potential interactions between cell populations from different lineages, we generated an *in vivo* atlas following the same steps as for the epithelial atlas described above, but without the subsetting to epithelial cells. In addition, no clustering was performed to assign cells to cell types. Instead, we proceeded in a stepwise manner. First, all cells that were part of the epithelial atlas were assigned the cell type label from there. Next, all remaining cells that were also part of the HECANext, all remaining cells that were also part of the HECA were assigned the label from there (immune cells were assigned the fine-grained labels from the dedicated analysis of the immune lineage)^31^. Finally, we used SingleR to predict labels for all remining cells, using the labelled cells as the reference data set.

Next, we subset the complete atlas to only samples from the menstrual and proliferative stages, and ran CellChat (v1.6.1) on this subset, following the workflow outlined in the package vignette^87^. Cell types with less than 100 cells were excluded, and only ‘Secreted Signaling’ interactions were included.

### Generation of WNT7A knock-out organoid lines

EOs were collected and processed into single cell suspension as described in previous section. Nucleofection was performed with the D-Nucleofector (Lonza Bioscience, #AAF-1003X) using the AMAXA P3 Primary Cell 4D-NucleofectorTM X Kit S (Lonza Bioscience, V4XP-3032). After centrifugation (6 minutes at 0.6 rcf), single cells were split in equal numbers per condition (∼500,000 cells) and diluted in 20 μl P3 buffer (3.6 μl supplement and 16.4 μl solution per condition). Alt-R® S.p. HiFi Cas9 Nuclease V3 (IDT Lubioscience, 1081060) was diluted in PBS to a final concentration of 24.4 μM. Multi-sgRNAs were diluted in duplex buffer to a final concentration of 44.4 μM. Equal volumes of Cas9 nuclease and sgRNAs were mixed for 20 min at room temperature for the formation of ribonucleoprotein (RNP) complex. Per condition, 20 μl of cells resuspended in P3 buffer were mixed with 2 μl of the RNP complex. For enhanced KO efficiency, 1 μl of a non-specific template (100 μM stock concentration) was used. Cells were then loaded on the cassette for nucleofection. Nucleofected cells were washed with EOM, centrifuged (6 min, 0.6 rcf) and resuspended in cold Matrigel (Corning, 356231). Sequences of sgRNAs and DNA template are listed in Supplementary Table 4. For generation of KO clones, single EOs were picked, broken both with automatic pipette and manually and finally plated in 5 μl Matrigel droplets in a 96-well plate. Individual clones were expanded for downstream analysis.

### RNA extraction

EOs were collected and Matrigel was removed using Cell Recovery Solution (Corning, 354253) for 50 min maximum on ice followed by one wash with cold PBS pH (7.4) (Gibco, 10010-015). Total RNA was extracted from organoid pellets using the RNeasy Micro kit with on column DNase treatment (Qiagen, 76004), following manufacturer’s instructions. The RNA was resuspended in 14 μl RNase-free water. The purity and concentration of the RNA was determined using the UV-Vis Spectrophotometer NP80 (Implen).

### cDNA synthesis

For cDNA synthesis, 500 ng to 1 μg of total RNA (depending on material availability) was reverse transcribed using SuperScript VILO cDNA Synthesis Kit (Thermo Fisher Scientific, 11754050) following manufacturer’s instructions. The extracted RNA was diluted into 5X VILO Reaction Mix containing random primers, dNTPs, and MgCl2 and 10X SuperScript III Enzyme Blend containing SuperScript III Reverse Transcriptase, RNase Recombinant Ribonuclease Inhibitor, and proprietary helper protein. The samples were then incubated for 10 minutes at 25°C, 1 hour at 42°C and 5 minutes at 85°C. An RNA sample without Reverse transcriptase was used as control for genomic DNA contamination.

### Real-time quantitative PCR (RT-qPCR)

RT-qPCR was performed with the StepOnePlus PCR system (Applied Biosystems) using TaqMan Fast Advanced Master Mix (Thermo Fisher Scientific, 4444557) and Taqman gene specific primer probes (*SCGB2A2*; Hs00935948, *FOXJ1*; Hs00230964, *MKI67*; Hs00606991, *WNT7A*; Hs00171699 (Thermo Fisher Scientific)), following manufacturer’s protocol. The cycling conditions followed were 20 seconds at 95°C and 40 cycles of 3 seconds at 95°C followed by 30 seconds at 60°C. Expression levels were calculated using the comparative Cycle threshold (Ct) method. The geometric means of *HPRT1*, *TOP1*, and *TBP* housekeeping genes were used for normalization of the relative gene expression levels. Normalized expression levels were calculated as 2^-ΔCt^ where ΔCt= Ct_gene of interest_ – Ct_geometric mean of housekeeping genes_. Each qRT-PCR reaction was performed in technical duplicates, and a non-template control was always included.

### Western Blotting

EOs were collected and Matrigel was removed using Cell Recovery Solution (Corning, 354253) for 50 min maximum on ice. EOs were then washed with cold PBS PH (7.4) (Gibco, 10010-015) and pellets were stored at -80°C. For lysis, pellets were resuspended in 500 μL RIPA lysis buffer (prepared in-house) supplemented with 1× Halt Protease Cocktail Inhibitor (Thermo Fisher Scientific, 1862209) and incubated on ice for 30 minutes. Lysates were centrifuged at 15,000×g for 20 minutes at 4°C, and the supernatant was collected for further analysis. Protein concentration was determined using the Pierce BCA Protein Assay Kit (Thermo Fisher Scientific, 23225) following manufacturer’s instructions. Protein lysates (∼10 μg) were diluted in 4× Laemmli SDS sample buffer (Thermo Fisher Scientific, J63615) containing 50 mM DTT (Thermo Fisher Scientific, D9779). Samples were boiled for 5 minutes at 95°C and loaded into SDS-PAGE gels alongside a molecular weight standard. Proteins were transferred to nitrocellulose membranes (Bio-Rad, 1704158) using the Turbo Transfer System (25 V, 1.3 A, 7 minutes). Membranes were blocked with 1× TBST containing 5% milk (Biotium, 22012) for 1 hour at room temperature. Membranes were incubated overnight at 4°C with primary antibody WNT7A (Abcam, ab274321) (1:1000) diluted in 1× TBST containing 5% milk. Tubulin (Thermo Fisher Scientific, t5168 (1:2000) was used as loading control. Goat Anti-Rabbit, (Bio-Rad, 1706515) secondary antibody conjugated to HRP was diluted 1:5000 in 1× TBST with 5% milk and incubated for 45 minutes at room temperature. Membranes were washed twice with 1× TBST and developed using Amersham ECL Select™ detection reagent (Cytiva, RPN2235). Chemiluminescence was detected using the Amersham imaging system.

### ELISA

Supernatants of EOs in the IVMC protocol and control EOs (B_NH) at Hormonal Withdrawal 48h, Post-breakdown 24h, E2-diff 48h were collected and stored at -80°C. Protein specific ELISA kits were used for the detection of CXCL8/IL-8 (Invitrogen, KHC0081), following manufacturer’s instructions. The calculated protein concentrations (pg/ml) were normalized for organoid confluency using the RNA concentration per well. Statistical analysis was performed using two-way ANOVA (hormonal treatment × timepoint), with Tukey’s multiple comparisons test.

## Abbreviations

ALDH1A1: aldehyde dehydrogenase 1 A1
B_NH: breakdown_no hormones
cAMP: cyclic AMP
CPM: counts per million
DEA: differential expression analysis
DEG: differentially expressed genes
E2: estrogen
ECM: Extracellular matrix
EOs: endometrial organoids
EOM: endometrial organoid medium
ESR1: estrogen receptor 1 (gene)
FUT4: fucosyltransferase 4
FZD: frizzled
IHC: immunohistochemistry
IVMC: *in vitro* menstrual cycle
KO: knock-out
KRT13: keratin 13
LGR5: leucine-rich repeat-containing G-protein-coupled receptor 5
NB_H: no breakdown_no hormones
NB_NH: no breakdown_no hormones
P4: progesterone
PAEP: progestagen associated endometrial protein
PCA: principal component analysis
PGR: progesterone receptor (gene)
PR: progesterone receptor (protein)
PLAUR: plasminogen activator urokinase receptor
PRL: prolactin
qRT-PCR: quantitative real-time PCR
RNAseq: RNA sequencing
scRNAseq: single cell RNA sequencing
sgRNA: single guide RNA
SOX9: SRY-Box Transcription Factor 9
SSEA-1: stage specific embryonic antigen-1
THBS1: thrombospondin
TPM: transcripts per million
WNT7A: Wingless-Type
MMTV: Integration Site Family, Member 7A
WT: wild type

## Code availability

All source code used for analysis and generation of plots, as well as additional supplementary tables, are available from GitHub: https://github.com/fmicompbio/IVMC-protocol.

## Acknowledgements

We are grateful to A. Moffett, P. Liberali and C. Tsiairis for valuable feedback on the manuscript. We are grateful to the Turco lab: U. Kilik, V. Bondarenko, K. Guja Jarosz, G. Guntri and particularly E. Magistrati and I. Calvi, for their support and contribution to discussions. We would also like to thank M. Huch and H. Critchley for their critical feedback on the project and D. Schübeler and H. Grosshans for their advice and support. Thank you to members of the Facility for Advanced Imaging and Microscopy, L. Gelman, L. Plantard, M. Bourbon, J. Eglinger and T.-O. Buchholz for their technical support with imaging and signal quantifications. We are grateful to the functional genomics facility, S. Smallwood and S. Aluri for coordinating and performing RNA sequencing experiments. We would like to thank T. Cindrova-Davies, G. J. Burton, K. Elder and Bourn Hall Fertility Clinic for the derivation of endometrial organoid cultures. This work was supported by the Novartis Research Foundation.

## Author contributions

K.N and M.Y.T. designed experiments. K.N., W.Y., L.K. performed experiments. K.N, S.C, H.R.H. analysed data. K.N and M.Y.T. interpreted the data. K.N. and M.Y.T. wrote the manuscript. K.N, W.Y., L.F., L.K. S.C, H.R.H., M.Y.T. reviewed the manuscript.

---

**Extended Data Figure 1:**
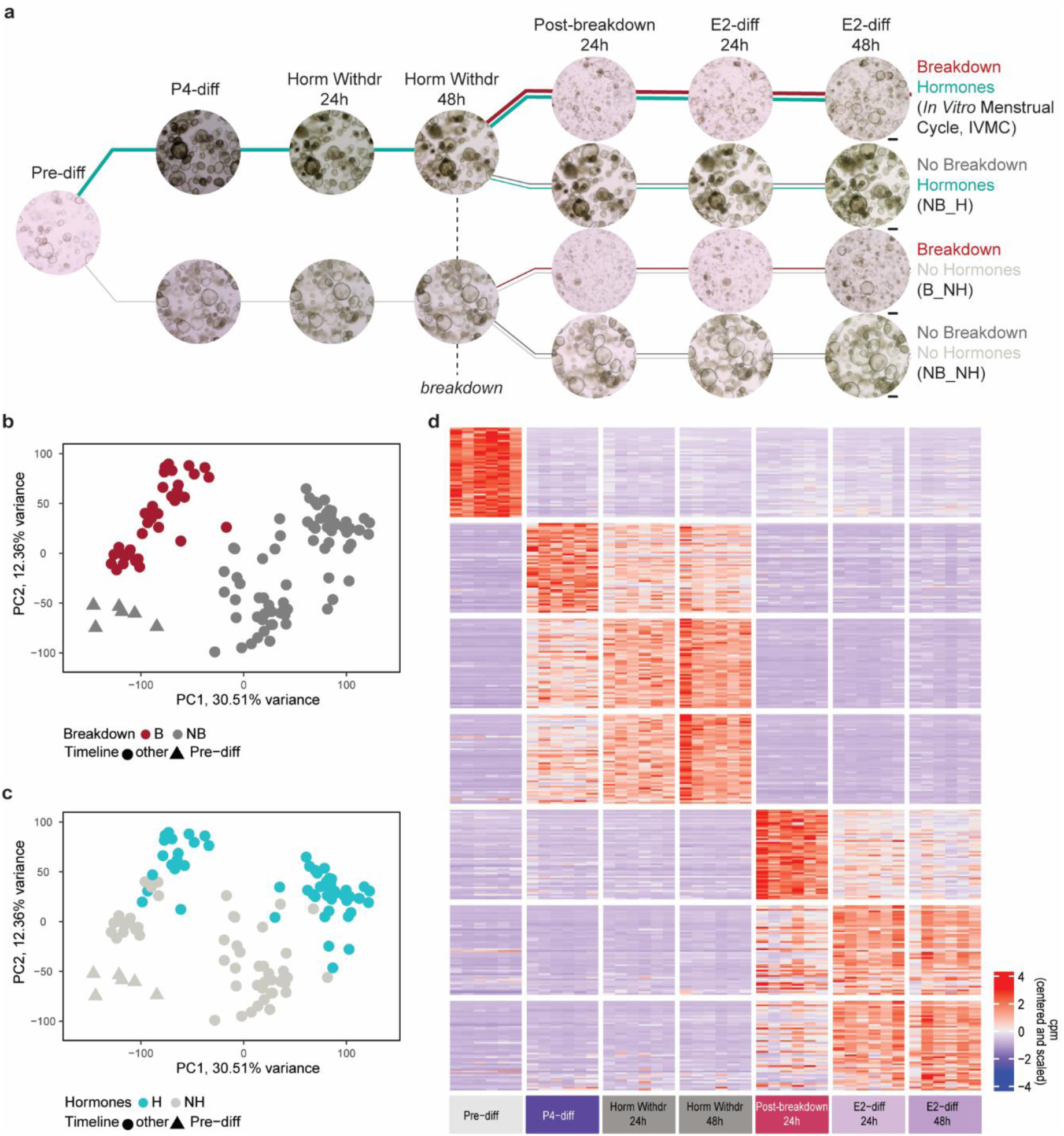
Bulk RNAseq of EOs undergoing the IVMC protocol. (**a**) Timeline of collection of EOs for bulk RNAseq (n=6 independent EO lines). The thicker lines represent the IVMC protocol. Control conditions include: EOs not broken down and treated with hormones (No Breakdown Hormones; NB_H); EOs broken down but not treated with hormones (Breakdown No Hormones; B_NH); or EOs neither broken down nor treated with hormones (No Breakdown No Hormones; NB_NH). Representative brightfield images for each timepoint are shown. Scale bar, 200 μm. (**b**) PCA of batch corrected samples from all conditions coloured by the breakdown variable. (**c**) PCA of batch corrected samples from all conditions coloured by the hormonal treatment variable. (**d**) Heatmap displaying the top 50 differentially upregulated genes from comparisons of each timepoint against all other timepoints using batch corrected samples from the IVMC protocol.

**Extended Data Figure 2:**
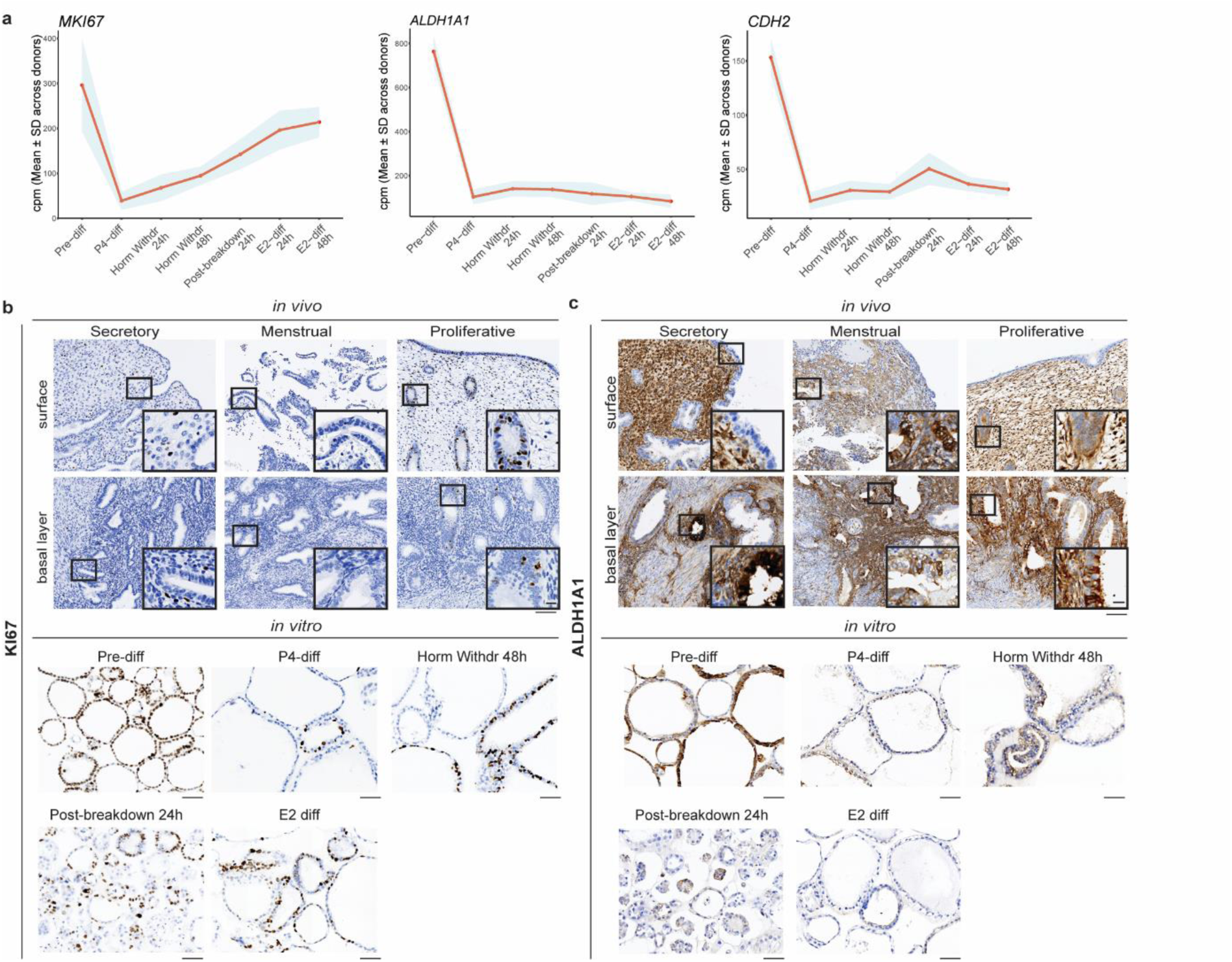
Expression of Pre-diff markers in EOs undergoing the IVMC protocol. (**a**) Line plots showing the mean expression levels in cpm of *MKI67*, *ALDH1A1*, and *CDH2* across the IVMC protocol. The red line represents the mean cpm values across donors (n=6 independent EO lines), while the shaded blue region indicates the standard deviation around the mean. (**b, c**) Representative IHC images for KI67 (b) and ALDH1A1 (c) in sections from secretory, menstrual, and proliferative endometrium (n=3 donors for each phase) and EOs across the IVMC protocol (n=4 independent EO lines). Black boxes indicate area shown at higher magnification (inset). Scale bars of tissue sections, 100 μm (main), 15 μm (inset). Scale bars of EO sections, 50 μm.

**Extended Data Figure 3:**
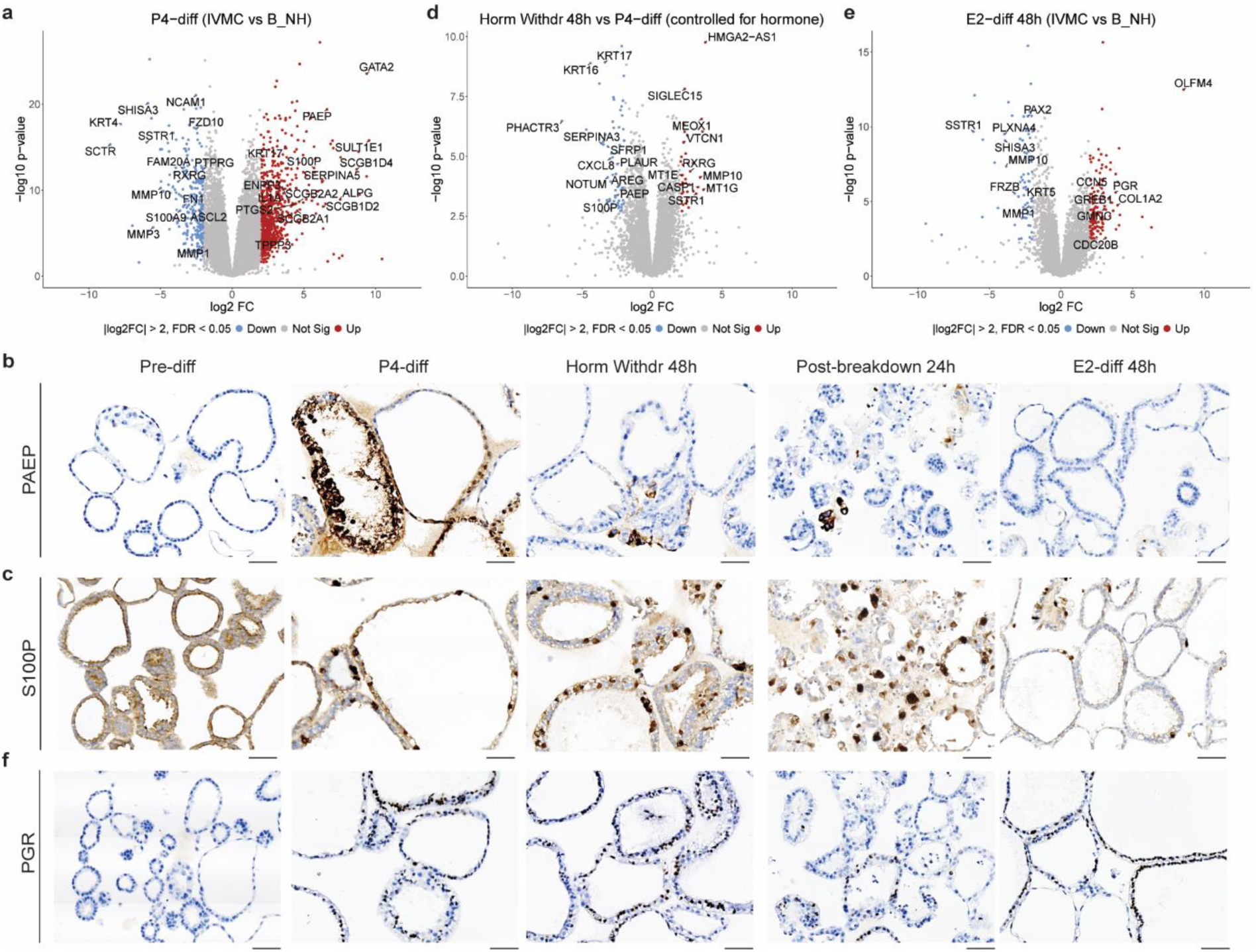
The effect of hormonal treatment in EOs undergoing the IVMC protocol. (**a**) Volcano plot highlighting selected DEGs comparing EOs at P4-diff phase of the IVMC protocol to control EOs (B_NH) at the same timepoint. (**b, c**) Representative IHC images for PAEP (b) and S100P (c) in sections from EOs across the IVMC protocol (n=4 independent EO lines). Scale bar, 50 μm. (**d**) Volcano plot highlighting selected DEGs comparing EOs of the IVMC protocol at P4-diff phase to EOs at Horm Withdr 48h corrected for control (NB_NH) conditions. (**e**) Volcano plot highlighting selected DEGs comparing EOs between E2-diff 48h phase of the IVMC protocol to control EOs (B_NH) at the same timepoint. (**f**) Representative IHC images for PGR in sections from EOs across the IVMC protocol (n=4 independent EO lines). Scale bar, 50 μm. Abbreviations: B_NH; Breakdown No Hormones, NB_NH; No Breakdown No Hormones.

**Extended Data Figure 4:**
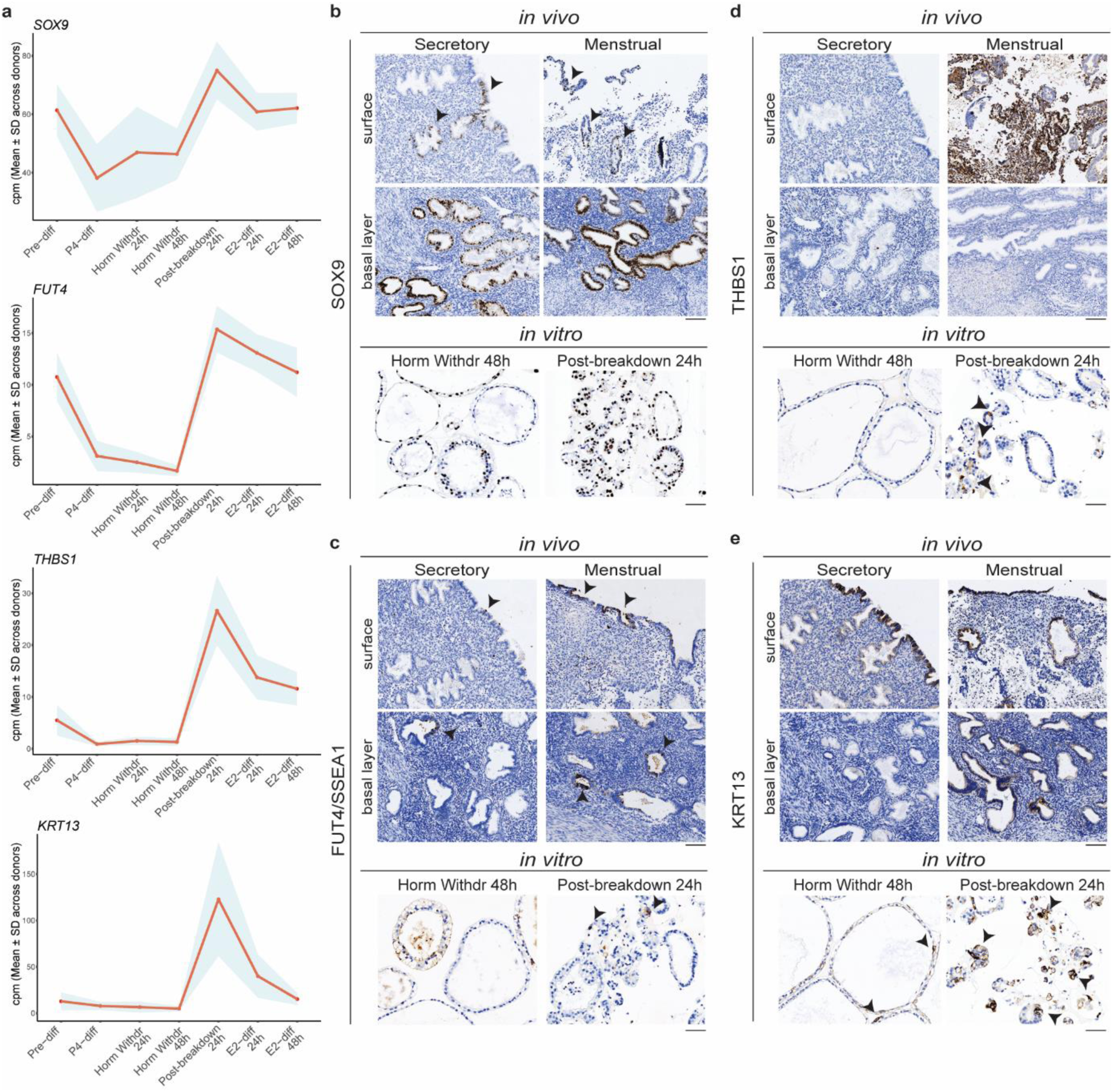
The effect of mechanical breakdown in EOs undergoing the IVMC protocol. (**a**) Line plots showing the mean expression levels in cpm of *SOX9*, *FUT4*, *THBS1* and *KRT13* genes across the IVMC protocol. The red line represents the mean cpm values across donors (n=6 independent EO lines), while the shaded blue region indicates the standard deviation around the mean. (**b-e**) Representative IHC image for SOX9 (b), SSEA1 (c), THBS1 (d), and KRT13 (e) staining in sections from functional and basal layers of secretory and menstrual endometrium (n=3 donors for each phase) and EOs at Horm Withdr 48h and Post-breakdown 24h (n=4 independent EO lines). Black arrowheads indicate positive cells. Scale bars of tissue sections, 100 μm. Scale bars of EO sections, 50 μm.

**Extended Data Figure 5:**
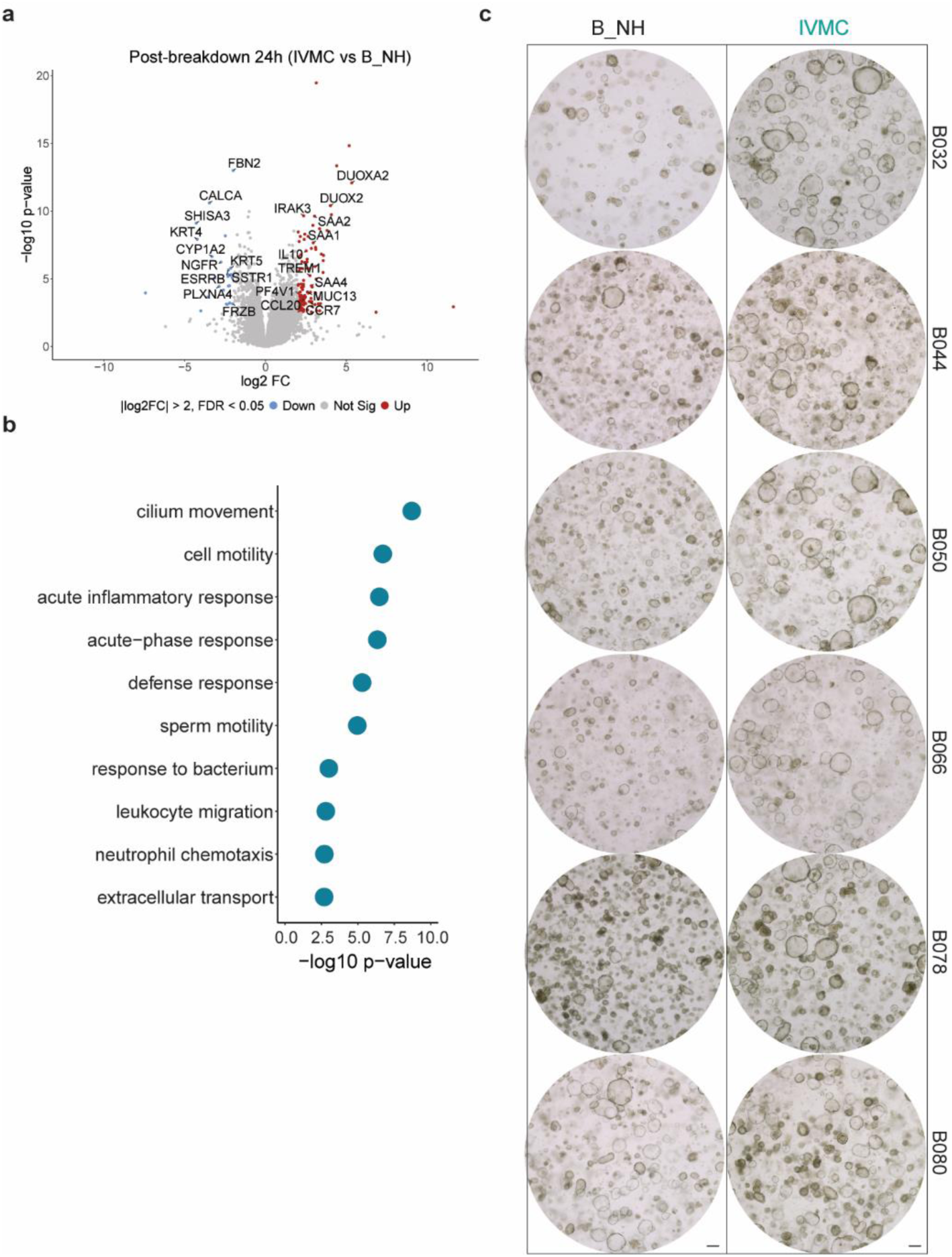
The effect of hormonal treatment in the response of EOs to breakdown. (**a**) Volcano plot highlighting selected DEGs comparing EOs at Post-breakdown 24h phase of the IVMC protocol to control EOs (B_NH) at the same timepoint. (**b**) Biological processes enriched in Post-breakdown 24h phase of the IVMC protocol using genes upregulated in (a). (**c**) Brightfield images from 6 independent EO lines at E2-diff 48h phase of the IVMC protocol and control EOs (B_NH) at the same timepoint. Scale bar, 200 μm. Abbreviations: B_NH; Breakdown No Hormones.

**Extended Data Figure 6:**
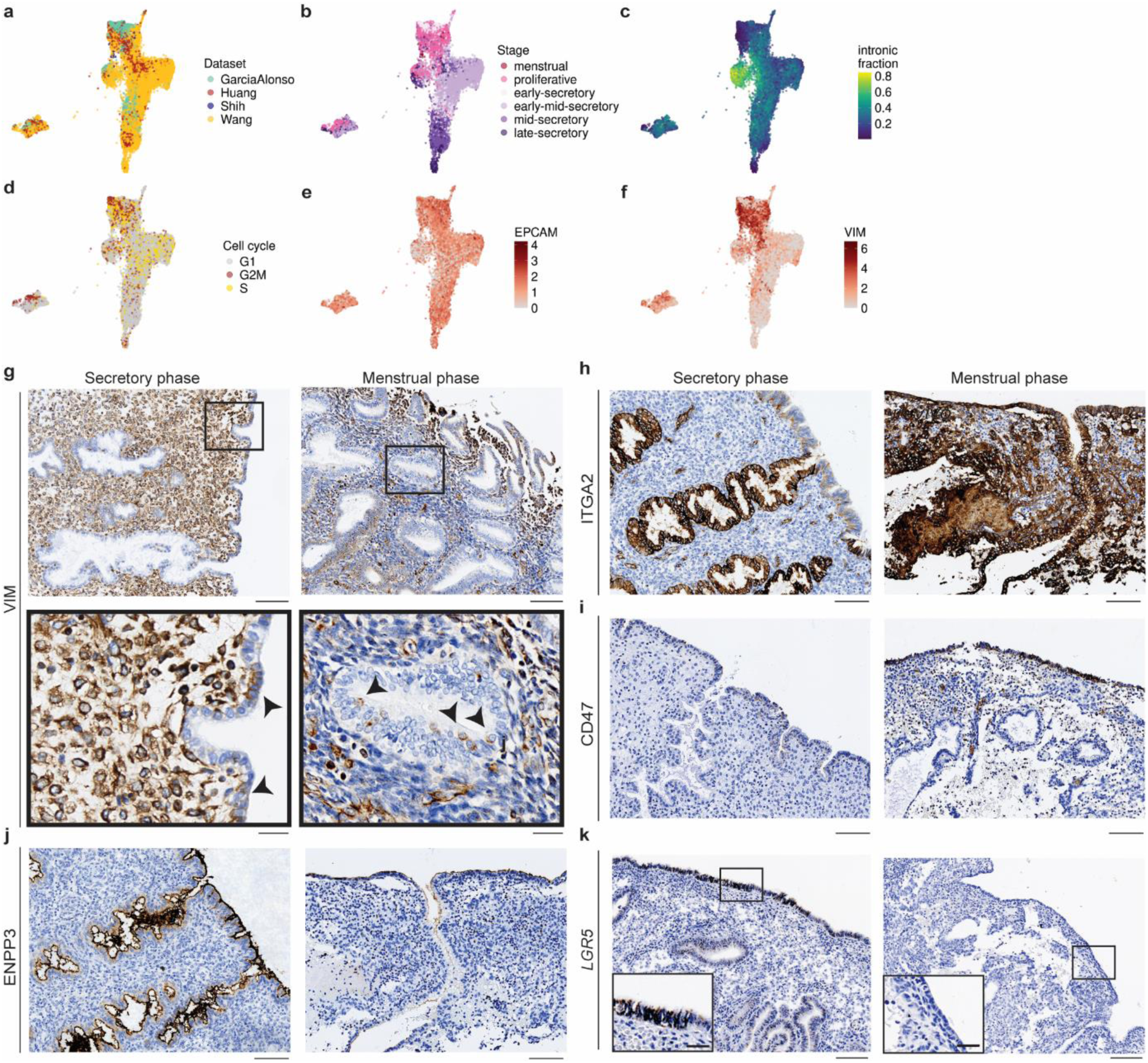
Description of epithelial cell populations *in vivo*. (**a-d**) UMAP visualization of epithelial cells from scRNAseq data of endometrial tissue coloured by the integrated datasets (a), stage of the menstrual cycle (b), intronic fraction (c), cell cycle phase (d). (**e-f**) UMAP visualization showing the log₂-transformed expression of *EPCAM* and *VIM* aross individual cells *in vivo*. (**g**) Representative IHC image for VIM in sections secretory and menstrual phase endometrium (n=3 donors for each phase). Scale bars, 100 μm (main), 20 μm (inset). (**h, i, j**) Representative IHC images for ITGA2, CD47 and ENPP3 in sections from secretory and menstrual phase endometrium (n=3 donors for each phase). Scale bars, 100 μm. (**k**) *In situ* hybridization showing localization of *LGR5* transcripts in secretory and menstrual phase endometrium (n=1 donor for each phase). Scale bars, 100 μm (main), 50 μm (inset).

**Extended Data Figure 7:**
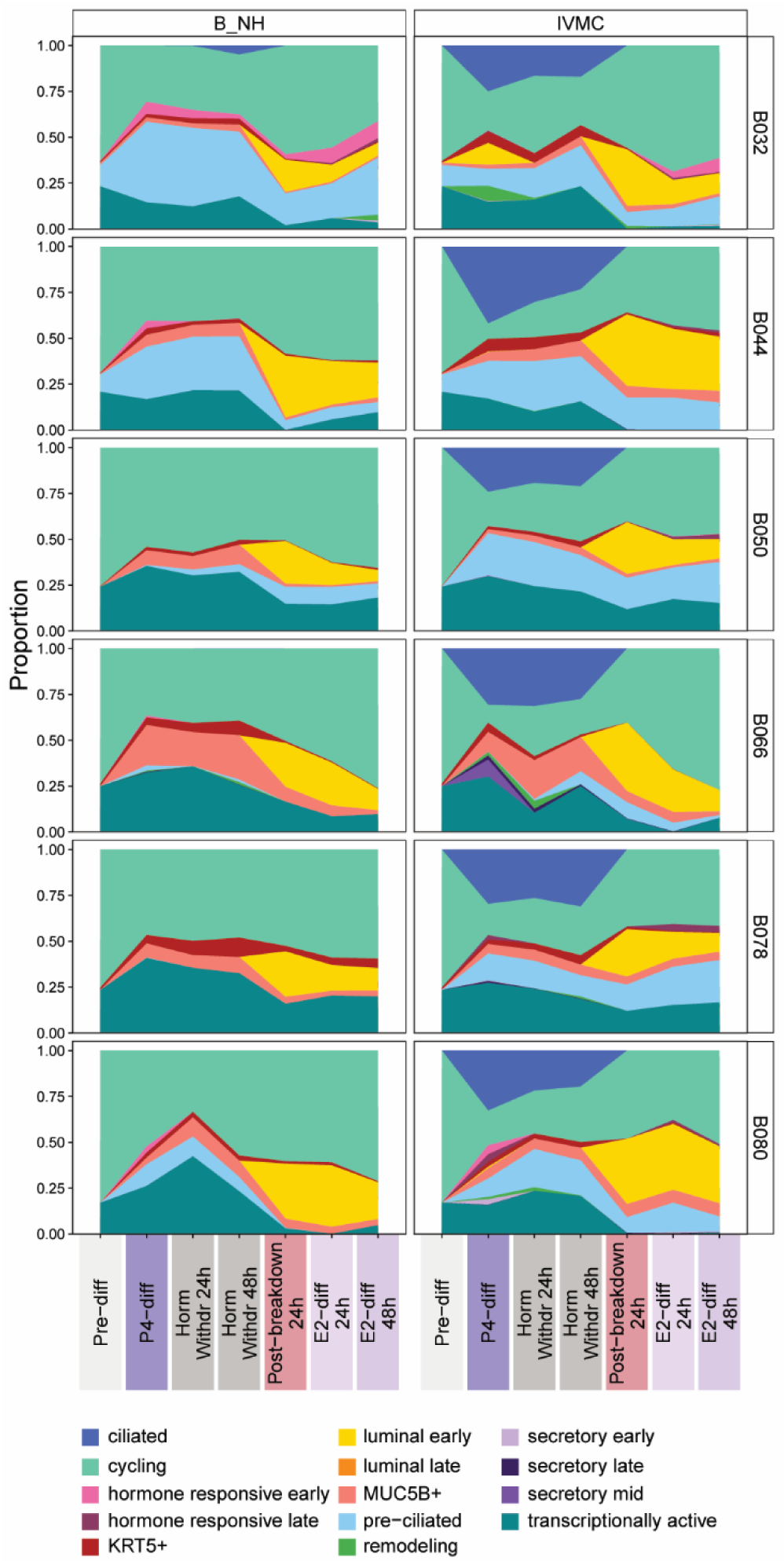
Deconvolution of expression profiles of EOs undergoing the IVMC protocol into *in vivo* epithelial cell populations. Area plots showing the relative abundance of the deconvolved *in vivo* epithelial cell clusters represented in EOs from all donors across the IVMC protocol and control EOs (B_NH). Abbreviations: B_NH; Breakdown No Hormones.

**Extended Data Figure 8:**
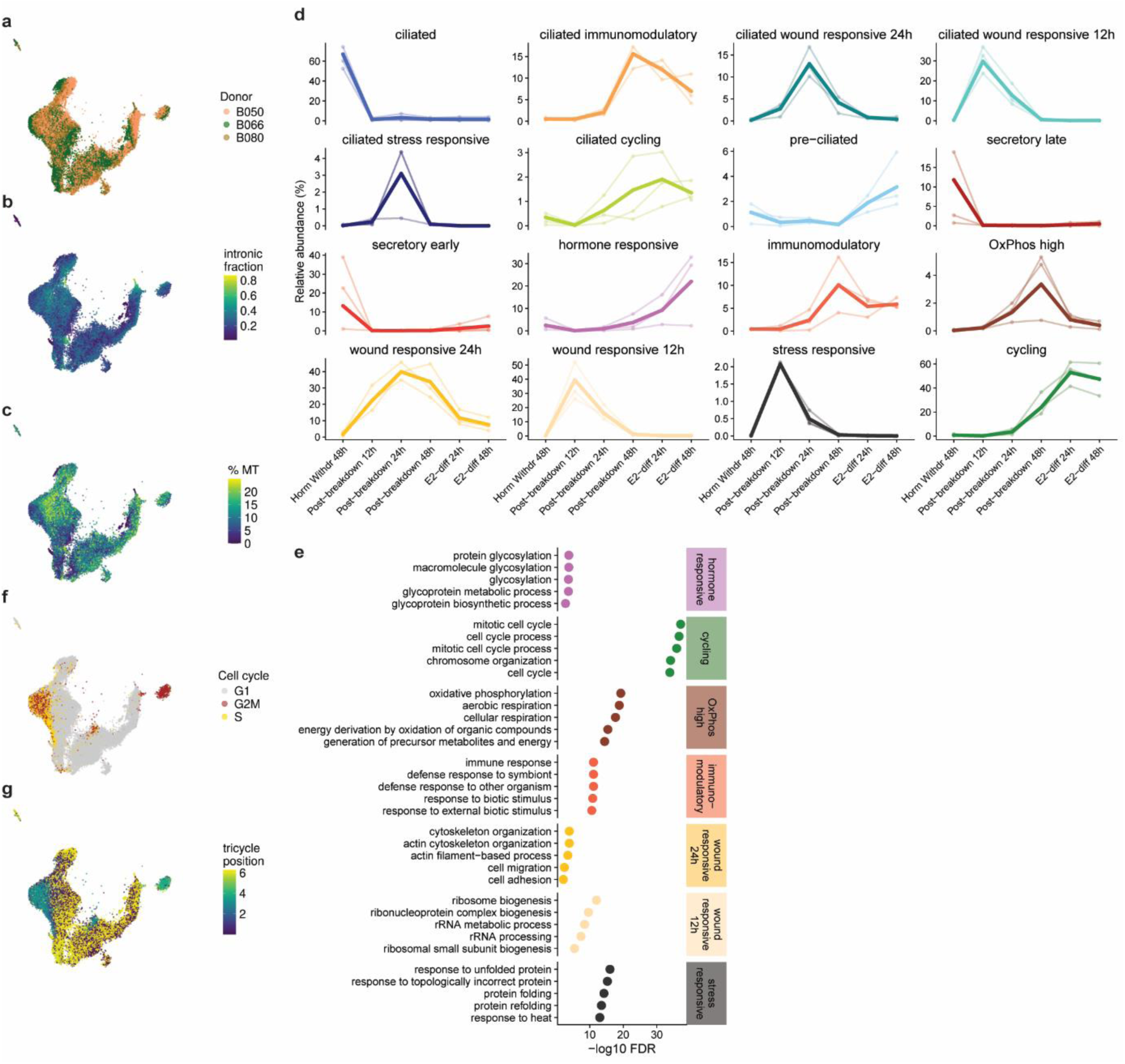
ScRNAseq analysis of EOs subjected to the IVMC protocol. (**a-c**) UMAP visualizations of epithelial cells in the IVMC protocol coloured for independent EO lines (a), intronic fraction (b) and percentage of counts coming from mitochondrial genes (c). (**d**) Relative abundance plots of all cell clusters across the different timepoints of the IVMC protocol. (**e**) Biological processes enriched in non-ciliated populations: stress responsive, wound responsive 12h, wound responsive 24h, immunomodulatory, OxPhos high, cycling and hormone responsive cell clusters. (**f, g**) UMAP visualizations of epithelial cells in the IVMC protocol coloured for the cell cycle stage (f) and tricycle position (g).

**Extended Data Figure 9:**
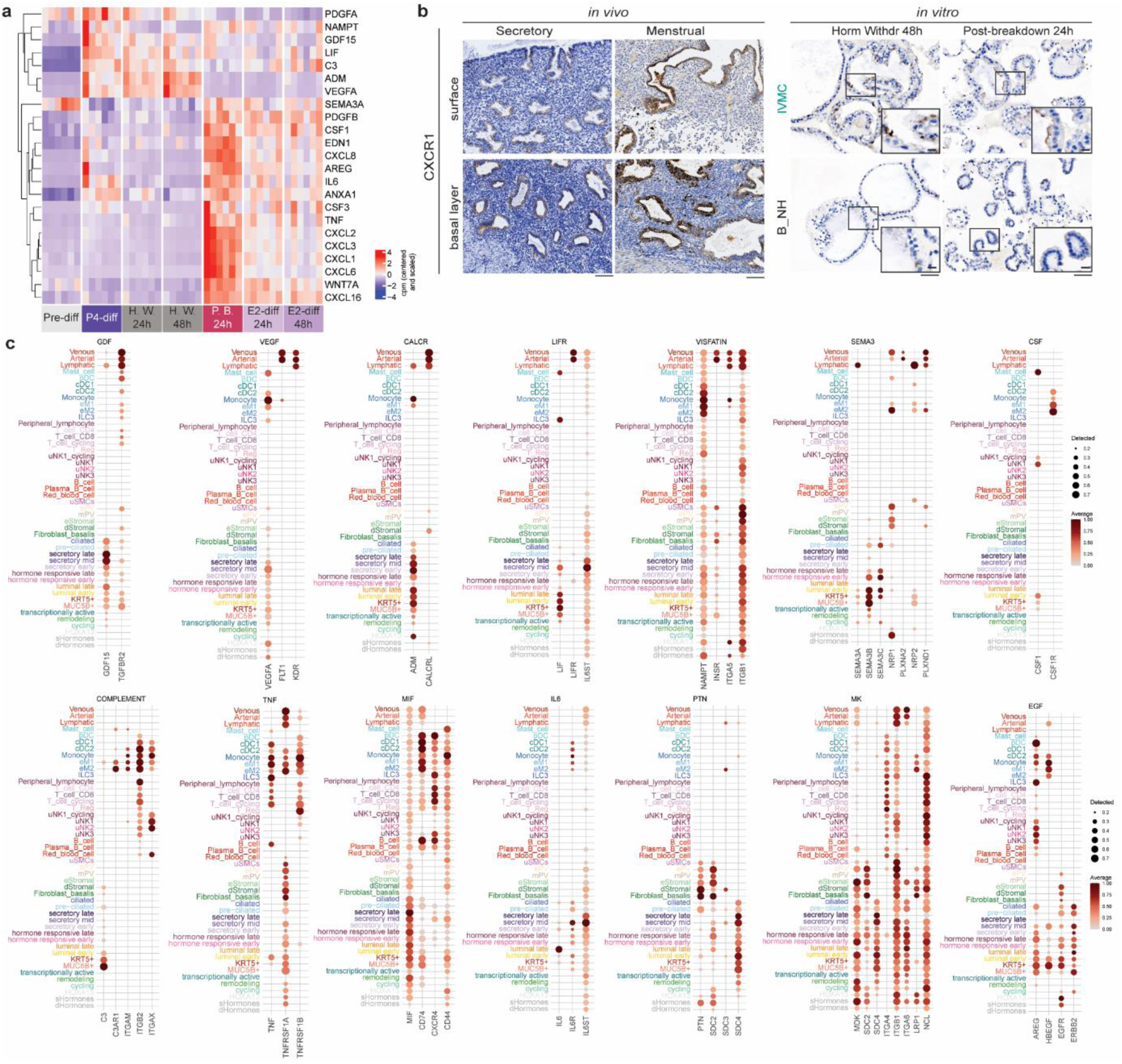
Expression of components of the signalling networks involved in interactions between luminal early epithelial cells and other cell types. (**a**) Heatmap depicting centered and scaled cpm of ligands involved in interactions between luminal early epithelial cells and other cell types, across batch corrected samples of EOs in the IVMC protocol. (**b**) Representative IHC image for CXCR1 (IL-8 receptor) staining in sections from functional and basal layers of secretory and menstrual endometrium (n=3 donors) and EOs at Horm Withdr 48h and Post-breakdown 24h (n=4 independent EO lines). Scale bars of tissue sections, 100 μm. Scale bars of EO sections, 50 μm, insets 25 μm. (**c**) Dot plot illustrating the expression of ligands and receptors of various signalling networks involved in interactions between luminal early epithelial cells and all the other cell types. Dot size represents the proportion of expressing cells, while colour denotes log2-transformed expression levels, normalised across all cell populations.

**Extended Data Figure 10:**
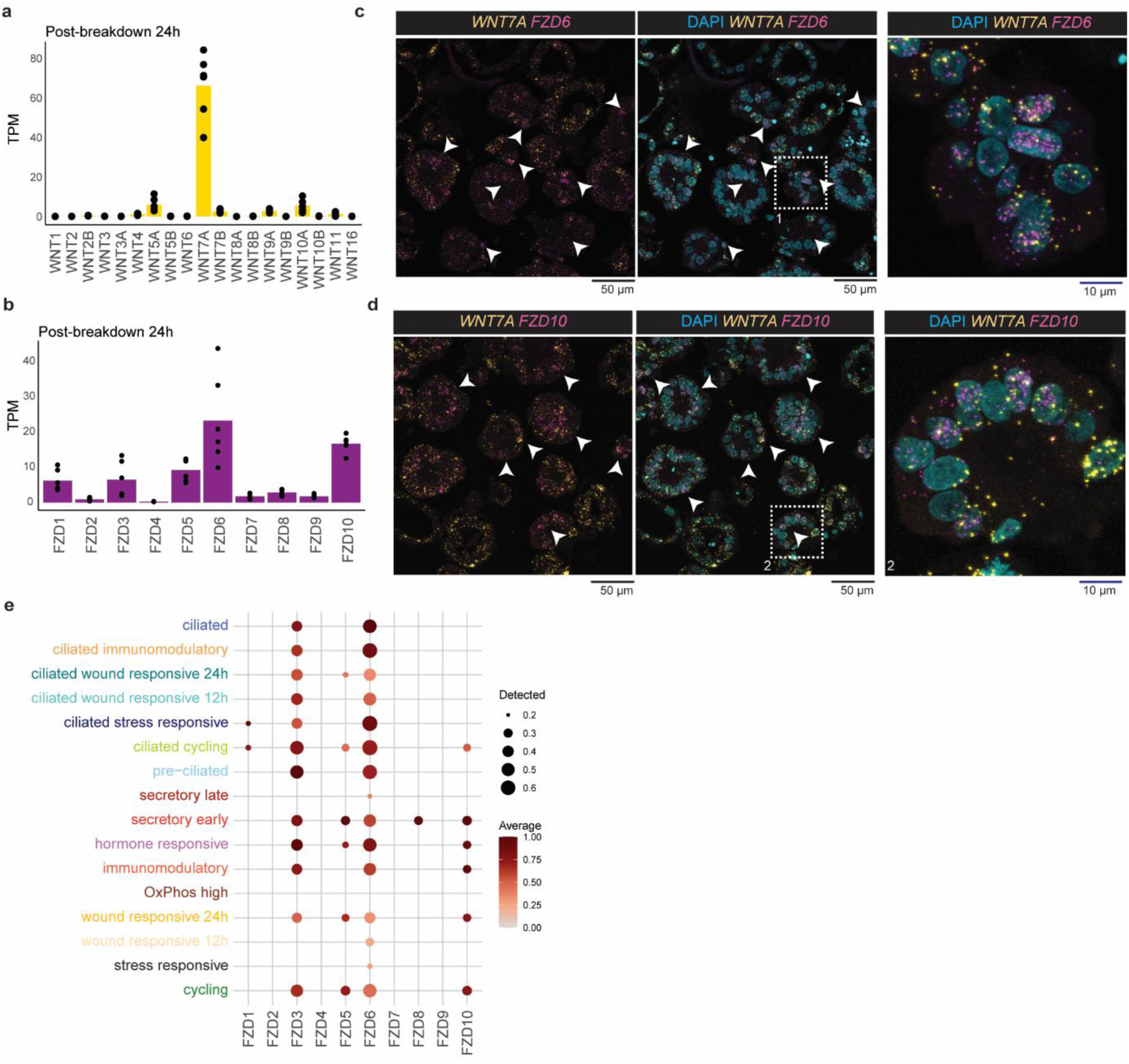
Expression of ligands and receptors of the WNT signalling pathway in EOs across the IVMC protocol. (**a, b**) Bar plot showing the average transcripts per million (TPM) of *WNT* ligands and *FZD* receptors at Post-breakdown 24h in EOs undergoing the IVMC protocol analysed with bulk RNAseq. (**c, d**) Multiplex ISH for *WNT7A* (yellow) together with *FZD6* or *FZD10* (in magenta) in sections of EOs at Post-breakdown 24h. White arrowheads point towards clear pattern of signal in individual nuclei (cyan). (**e**) Dot plot illustrating the expression of *FZD* receptors characteristic in each epithelial cell cluster in EOs subjected to the IVMC protocol. Dot size represents the proportion of expressing cells, while colour denotes log2-transformed expression levels, normalised across all cell populations.

**Supplementary Table 1.**
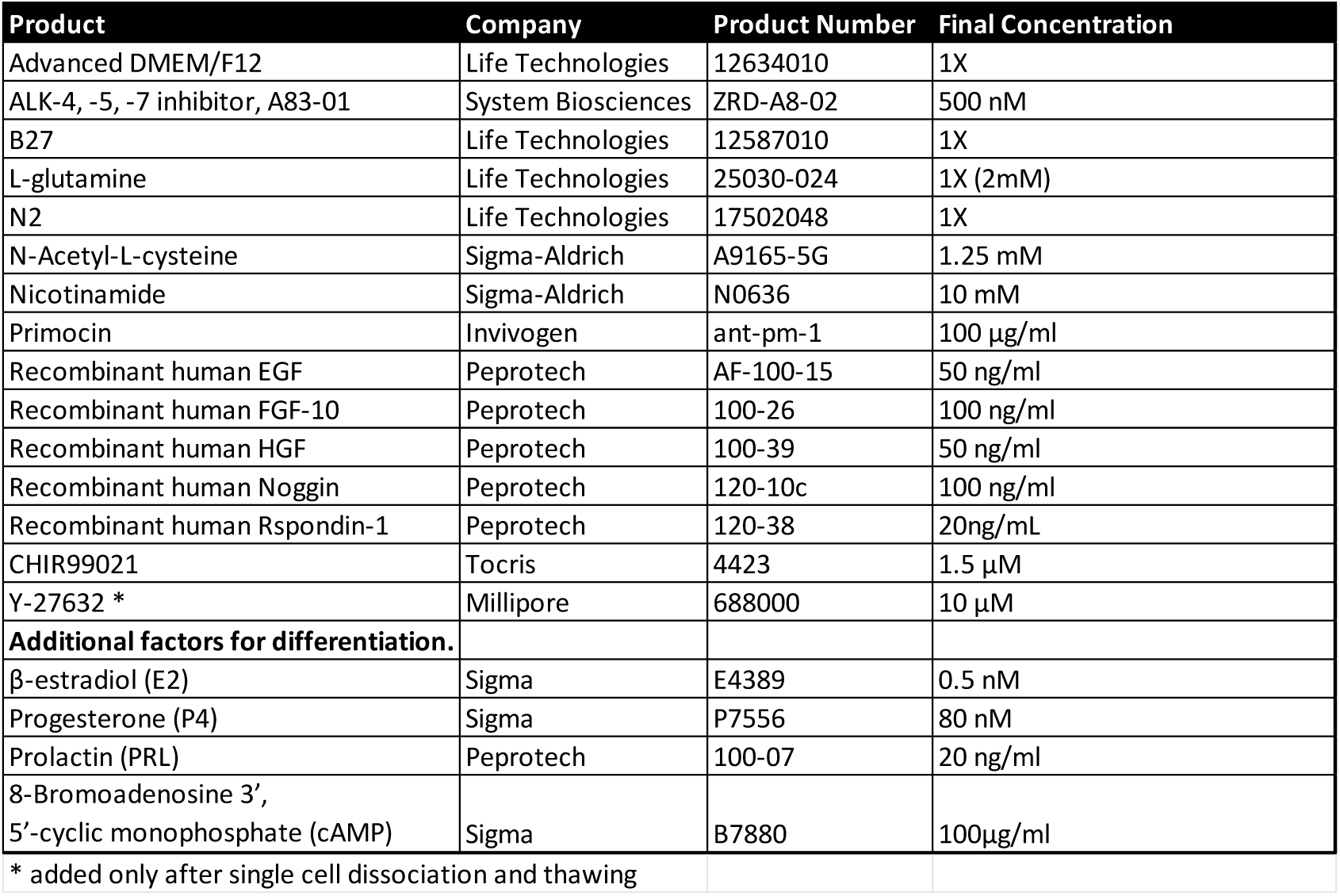
Composition of endometrial organoid medium (EOM) and differentiation medium.

| Product | Company | Product Number | Final Concentration |
| --- | --- | --- | --- |
| Advanced DMEM/F12 | Life Technologies | 12634010 | 1X |
| ALK-4, -5, -7 inhibitor, A83-01 | System Biosciences | ZRD-A8-02 | 500 nM |
| B27 | Life Technologies | 12587010 | 1X |
| L-glutamine | Life Technologies | 25030-024 | 1X (2mM) |
| N2 | Life Technologies | 17502048 | 1X |
| N-Acetyl-L-cysteine | Sigma-Aldrich | A9165-5G | 1.25 mM |
| Nicotinamide | Sigma-Aldrich | N0636 | 10 mM |
| Primocin | Invivogen | ant-pm-1 | 100 µg/ml |
| Recombinant human EGF | Peptotech | AF-100-15 | 50 ng/ml |
| Recombinant human FGF-10 | Peptotech | 100-26 | 100 ng/ml |
| Recombinant human HGF | Peptotech | 100-39 | 50 ng/ml |
| Recombinant human Noggin | Peptotech | 120-10c | 100 ng/ml |
| Recombinant human Rspodin-1 | Peptotech | 120-38 | 20ng/mL |
| CHIR99021 | Tocris | 4423 | 1.5 µM |
| Y-27632 * | Millipore | 688000 | 10 µM |
| **Additional factors for differentiation.** |  |  |  |
| β-estradiol (E2) | Sigma | E4389 | 0.5 nM |
| Progesterone (P4) | Sigma | P7556 | 80 nM |
| Prolactin (PRL) | Peptotech | 100-07 | 20 ng/ml |
| 8-Bromoadenosine 3',<br>5'-cyclic monophosphate (cAMP) | Sigma | B7880 | 100µg/ml |
| * added only after single cell dissociation and thawing |  |  |  |

**Supplementary Table 2.** Antibodies used for immunohistochemistry.

| Antibody | Product Number | Company | Species | Clonality | Antigen retrieval | Dilution |
| --- | --- | --- | --- | --- | --- | --- |
| ALDH1A1 | 36671 | Cell Signalling | Rabbit | monoclonal D9J7R | pH9 | 1:500 tissues/1:5000 organoids |
| ANXA1 | 32934 | Cell Signaling | Rabbit | monoclonal D5V2T | pH6 | 1:300 |
| CD47 | ab218810 | Abcam | Rabbit | monoclonal EPR21794 | pH9 | 1:500 |
| CK13 | ab92551 | Abcam | Rabbit | polyclonal | pH9 | 1:100 |
| CXCR1 (C-terminus) | LS-A808 | LifeSpan Bio | Rabbit | polyclonal | pH9 | 1:1000 |
| ENPP3 | HPA043772 | Atlas antibodies | Rabbit | polyclonal | pH9 | 1:600 |
| ITGA2/49b | NBP3-15644 | Novus | Rabbit | monoclonal 7O9H6 | pH9 | 1:600 |
| Ki67 | MA5-14520 | Invitrogen | Rabbit | monoclonal SP6 | pH9 | 1:400 tissues/1:1500 organoids |
| PAEP | ab270454 | Abcam | Rabbit | monoclonal EPR23251-87 | pH9 | 1:7000 tissues/1:750 organoids |
| PGR | 8757 | Cell Signalling | Rabbit | monoclonal D8Q2J | pH9 | 1:1000 tissues/1:300 organoids |
| PLAUR | ab218106 | Abcam | Rabbit | monoclonal EPR20191 | pH9 | 1:200 |
| S100P | ab124743 | Abcam | Rabbit | monoclonal EPR6142 | pH9 | 1:300 tissues/1:1500 organoids |
| SOX9 | ab185230 | Abcam | Rabbit | monoclonal EPR14335 | pH6 | 1:1000 |
| SSEA1 | ab218403 | Abcam | Rabbit | monoclonal 1478R | pH9 | 1:40 |
| THBS1 | RAB01344 | Abnova | Rabbit | monoclonal R06-5F3 | pH9 | 1:500 |
| VIMENTIN | 5741 | Cell Signaling | Rabbit | monoclonal D21H3 | pH6 | 1:750 |

**Supplementary Table 3.** Probes used for in situ hybridization.

| Gene | Product Number | Company |
| --- | --- | --- |
| Hs-FZD10-C3 | 477131-C3 | ACD |
| Hs-FZD6-C1 | 460541 | ACD |
| Hs-FZD6-C2 | 460541-C2 | ACD |
| Hs-LGR5-C1 | 311021 | ACD |
| Hs-WNT7A-C1 | 408231 | ACD |
| Hs-WNT7A-C2 | 408231-C2 | ACD |
| Hs-PPIB-C1 (positive control) | 313901 | ACD |
| Hs-UBC-C1 (positive control) | 310041 | ACD |
| DapB-C1 (negative control) | 310043 | ACD |

**Supplementary Table 4.**
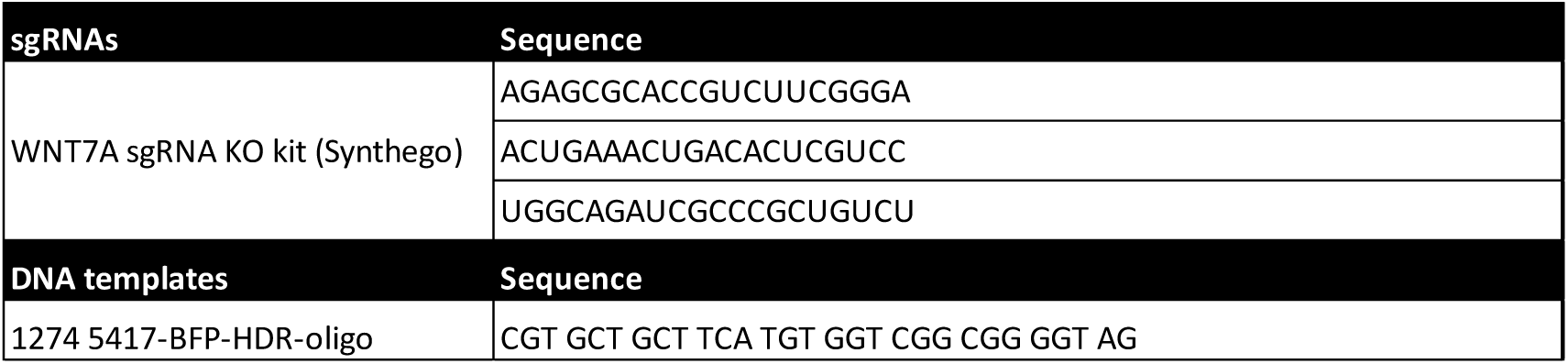
Sequences of sgRNAs and DNA templates used for CRISPR editing experiments.

